# Cell-type specific transcriptional networks in root xylem adjacent cell layers

**DOI:** 10.1101/2022.02.04.479129

**Authors:** Maria Amparo Asensi Fabado, Emily A Armstrong, Liam Walker, Giorgio Perrella, Graham Hamilton, Pawel Herzyk, Miriam L Gifford, Anna Amtmann

## Abstract

Transport of water, ions and signals from roots to leaves via the xylem vessels is essential for plant life and needs to be tightly regulated. The final composition of the transpiration stream before passage into the shoots is controlled by the xylem-adjacent cell layers, namely xylem parenchyma and pericycle, in the upper part of the root. To unravel regulatory networks in this strategically important location, we generated Arabidopsis lines expressing a nuclear tag under the control of the *HKT1* promoter. HKT1 retrieves sodium from the xylem to prevent toxic levels in the shoot, and this function depends on its specific expression in upper root xylem-adjacent tissues. Based on FACS RNA-sequencing and INTACT ChIP-sequencing, we identified the gene repertoire that is preferentially expressed in the tagged cell types and discovered transcription factors experiencing cell-type specific loss of H3K27me3 demethylation. For one of these, ZAT6, we show that H3K27me3-demethylase REF6 is required for de-repression. Analysis of *zat6* mutants revealed that ZAT6 activates a suite of cell-type specific downstream genes and restricts Na^+^ accumulation in the shoots. The combined Files open novel opportunities for ‘bottom-up’ causal dissection of cell-type specific regulatory networks that control root-to-shoot communication under environmental challenge.

## INTRODUCTION

Multicellular organisms are composed of different organs, tissues and cell types, which carry out distinct functions. In plants, the leaves are responsible for the assimilation of carbon while the roots take up water and mineral nutrients (Amtmann and Blatt, 2009). Within the roots, there is a further division of tasks along both the radial and the longitudinal axis (Barberon and Geldner, 2014). Transport of water and minerals occurs first in a radial manner from the epidermis towards the central cylinder (stele). Uptake into the xylem vessels then enables upwards longitudinal transport with the transpiration stream (Lucas et al., 2013). In the mature parts of the root system, the movement of substances into and out of the stele involves at least one trans-membrane step into the symplast because the apoplastic pathway is blocked by the Casparian strip and suberin barriers (Barberon and Geldner, 2014). A second trans-membrane step is required for loading and unloading of the (apoplastic) xylem vessels (Møller et al., 2009). It occurs either at the pericycle (Casimiro et al., 2003) or in the cells that are directly adjacent to the xylem, often called the xylem parenchyma (Maathuis et al., 2014) (Møller et al., 2009). Control of passage in these tissues is essential for regulating the content of the transpiration stream, which moves not only water and nutrients to the shoot, but also toxic substances (Mendoza-Cózatl et al., 2011; Munns and Tester, 2008) and signals (Notaguchi and Okamoto, 2015).

Specific gene expression in the root stele underpins important functions. For example, in *Arabidopsis thaliana*, several membrane transporters have been identified that either release or retrieve K^+^ or Na^+^ into and from the transpiration stream, including the K^+^ channel SKOR1 (Sharma et al., 2013), K^+^/H^+^ antiporter NPF7.3 (Li et al., 2017) and the Na^+^ transporters SOS1 (Shi et al., 2002) and HKT1 (Munns and Tester, 2008; Møller et al., 2009). All of them show preferential expression in root xylem-adjacent cells. Regulation of xylem content also relies on cell-type specific signals and signal perception. For example, induction of a NADPH oxidase in the root xylem parenchyma under salt stress leads to local generation of reactive oxygen species (ROS) and reduced root-to-shoot Na^+^ delivery (Jiang et al., 2013). The xylem Na^+^/K^+^ ratio is regulated by a radial ethylene signal from the root periphery to the stele, which indicates cell-type specific signal perception in the xylem-adjacent cell layers (Jiang et al., 2013). However, the individual components and targets of the regulatory network in this strategically important location remain to be elucidated.

To better understand the processes that control root-shoot communication we need to identify the genes expressed in the relevant cell types and we need to investigate their causal inter-relationships. FACS combined with microarray analysis or RNA-Sequencing (Birnbaum et al., 2003; Brady et al., 2007; Dinneny et al., 2008; Walker et al., 2017) and more recently with single-cell RNA-sequencing (Shulse et al., 2019; Jean-Baptiste et al., 2019, Wendrich et al., 2020, Long et al. 2021, Zhang et al 2021) have already delivered spatial expression maps of roots in Arabidopsis and rice. However, due to markers used, datasets for the stele are mostly limited to the young (meristematic) parts of the roots. Information on xylem-adjacent cells in mature root regions is currently based on computational inference rather than direct markers (Winter et al., 2007), although recent single-cell study on lateral-root forming regions in the differentiation zone has generated additional markers that could be used in the future (Gala et al. 2020).

Networks of genes co-expressed in the root stele also remain to be explored. Inference of cell-type specific gene regulatory networks from computational analysis of *cis*-motif enrichment (Jean-Baptiste et al., 2019; Sijacic et al., 2018) suggested that cell-type specific TFs extensively regulate each other (Sijacic et al., 2018), but some other mechanism must come into play to anchor this network in the specific location. Histone post-translational modifications are good candidates for a primary mechanism since they are important for transcriptional regulation during development (Roudier et al., 2011; Liu et al., 2010). H3K27me3 is a hallmark of gene repression (Zhang et al., 2007) and different H3K27me3 marking has been reported for individual plant cell types, such as root epidermal hair and non-hair cells (Deal and Henikoff, 2010), vascular cells (de Lucas et al., 2016) and guard cells (Lee et al., 2019). Moreover, several studies point to a role of H3K27me3 in establishing and maintaining cellular identity (Ikeuchi et al., 2015). However, recent study on guard cell lineages highlighted that the number of differentially H3K27me3-marked genes was small compared to the number of differentially expressed genes, and the authors proposed that epigenetic re-programming of a few core regulators could proliferate cell-type specificity into a large cell-type specific transcriptome (Lee et al. (2019).

Here we used a combination of FACS-RNA-seq and INTACT-ChIP-seq on *Arabidopsis thaliana* root samples to address the following questions: 1. Which genes are preferentially expressed in cell-types that control root xylem content? 2. What determines their cell-type specific expression? 3. Which core regulators control the cell-type specific transcriptional network?

Our results show that H3K27me3-mediated de-repression of a small set of transcription factors is sufficient for establishing large cell-type specific transcriptional networks, and we identify ZAT6 as one histone demethylation-dependent regulatory hub in root xylem-adjacent cell types.

## RESULTS

### Cell-tagging driven by the HKT1 promoter identifies co-expressed genes

For sorting of cell types controlling xylem content in the roots we took advantage of the cell-type specific expression and function of HKT1 (At4g10310), a plasma membrane transport protein that retrieves Na^+^ from the transpiration stream (Sunarpi et al., 2005, Møller et al., 2009). To enable analysis of the HKT1-expressing cell types by FACS and INTACT we generated a GATEWAY-destination vector allowing recombination of the HKT1-promoter sequence (Mäser et al., 2002) with the NTF cassette (Curtis and Grossniklaus, 2003), comprised of a nuclear envelope protein domain (WPP), green fluorescent protein (GFP) and a biotin acceptor peptide (BLRP), which acts as a substrate for *E*.*coli* biotin ligase (BirA). The resulting *pHKT1::NTF* construct was used to transform biotin-ligase expressing *A. thaliana* Col-0 *pACT2::BirA* plants (Deal and Henikoff, 2011). Stable homozygous lines carrying *pHKT1::NTF pACT2::BirA* (‘HKT1 INTACT lines’) produced a strong nuclear GFP signal in the cell layers surrounding the mature root xylem vessels, primarily xylem parenchyma and pericycle (Figure 1).

**Figure 1.**
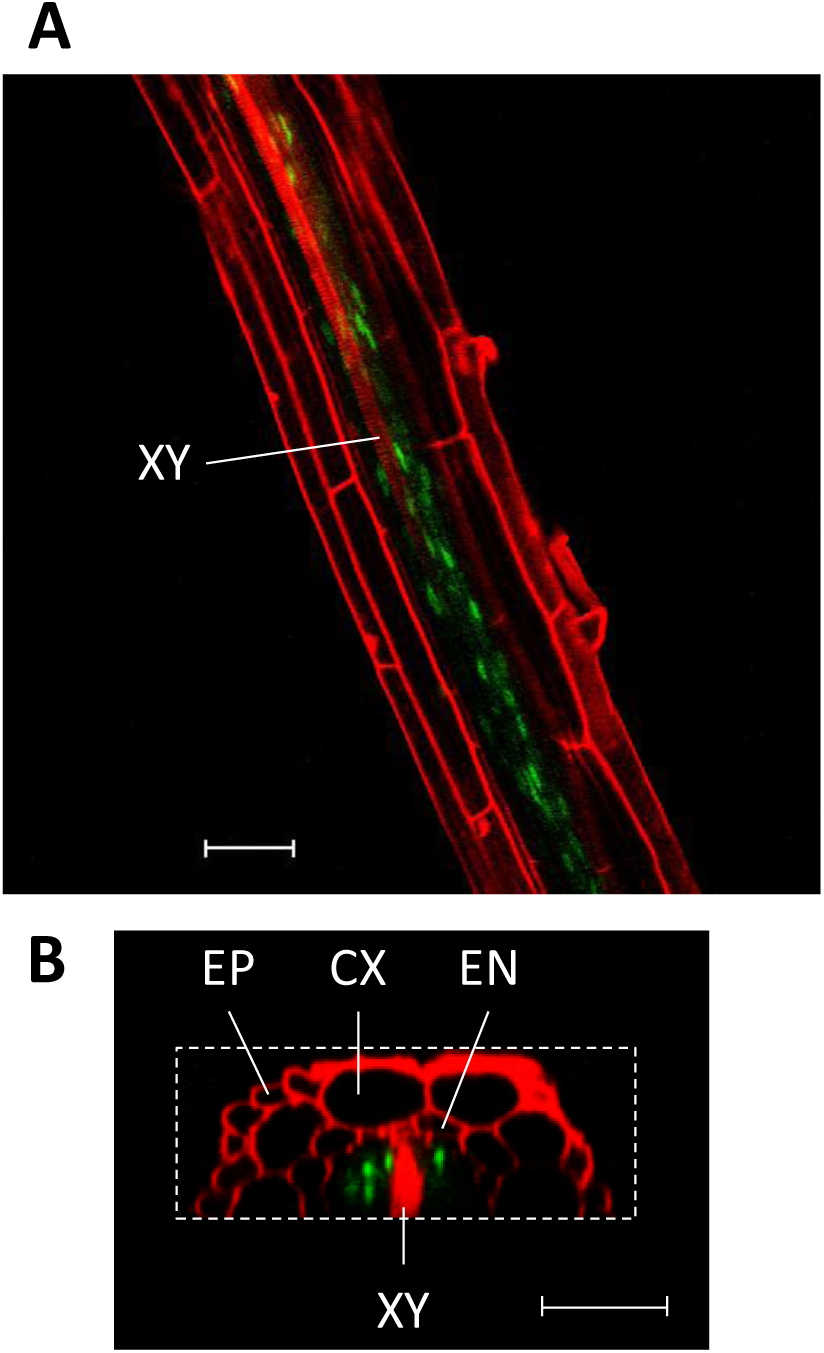
pHKT1-driven nuclear GFP-labelling of cell types surrounding the root xylem. GFP signal (green) in the root of *Arabidopsis thaliana* expressing *pHKT1::NTF* and *pACT2::BirA*. Cell walls were stained with propidium iodide (red) and images taken with the confocal microscope collecting fluorescent signals through 505-530 nm (GFP) and 560-615 (PI) nm filters after excitation at 488 nm. Z-stacks were taken at 1 μm intervals and combined with Image J software to construct the images. **A:** Longitudinal section through the centre of the primary root. **B:** Orthogonal projection representing a transverse cross section of the primary root, reconstructed from the Z-stacks. XY: xylem, EP: epidermis, CX: cortex, EN: endodermis. Scale bars: 50 μm.

To identify genes that are preferentially expressed in these cell types we harvested roots of 2-weeks old HKT1-INTACT plants from five independently grown batches, isolated protoplasts and carried out fluorescence-activated cell sorting (FACS). GFP-positive (‘tagged’) and GFP-negative (‘non-tagged’) cell samples were collected and subjected to RNA-sequencing. Principal component analysis (PCA) based on normalised mRNA levels in each gene showed a clear separation between the tagged and the non-tagged samples (PC1 explaining 77% of variation, Figure 2).

**Figure 2.**
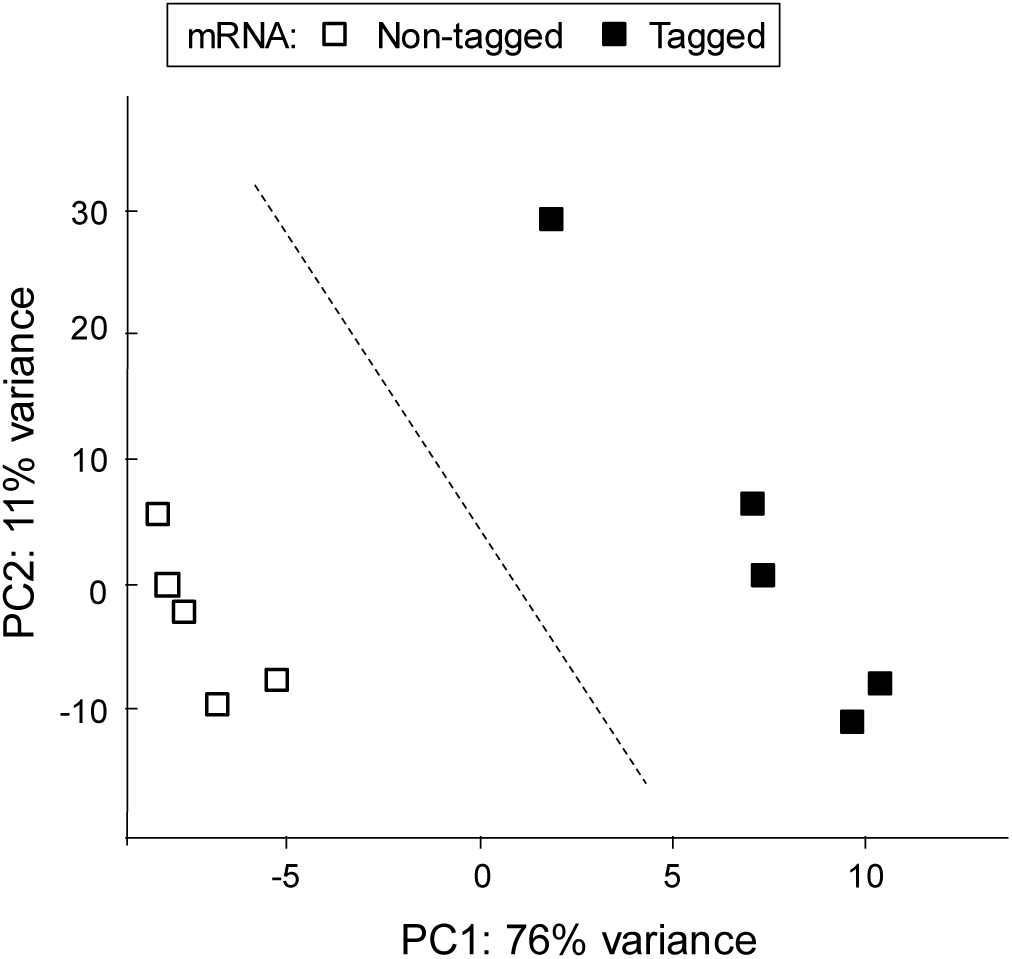
Principal component analysis of genome-wide transcript levels distinguishes between root cell types. Protoplasts were obtained from roots of *A. thaliana pHKT1::NTF pACT2::BirA* lines expressing GFP (as part of the NTF cassette) under the control of pHKT1, and subjected to Fluorescence Activated Cell sorting (FACS) followed by RNA-sequencing. Replicate samples were obtained from five independently grown plant batches. Black squares represent the pHKT1-NTF tagged protoplasts, open squares represent the samples containing the remaining (non-tagged) protoplasts. The PCA was based on normalised transcript levels for all genes. PC1 separates the tagged from the non-tagged samples explaining 76% of variation.

The highest statistically significant fold-change (tagged/non-tagged) was 101.5 (p = 3.26 × 10^−17^) for a gene encoding a glycosyl hydrolase (At1g58225). Preferential expression of HKT1 was 11.8-fold (p = 9.7 × 10^−8^). Other transporters with known specific expression in these cell-types also showed significantly higher transcript levels in the tagged samples than in the untagged, including SKOR (At3g02850, 5-fold, 10^−7^), which loads K^+^ into the xylem (Gaymard et al., 1998), and the nitrate transporters NRT1.5/NPF7.3 (At1g32450, 7-fold, 4×10^−10^) and NRT1.8/NPF7.2 (At4g21680, 11-fold, 10^−8^), which mediate xylem loading and unloading of nitrate, respectively (Li et al., 2017; Lin et al., 2008). Applying a threshold of at least 4-fold enrichment and p < 10^−4^ we identified 2020 genes with preferential expression in the tagged cells compared to the rest of the root, representing approximately 8% of the root expressed transcriptome (25144 genes, see Methods).

Analysis based on GO-terms and keywords using DAVID (Huang et al., 2009) revealed enrichment of annotations in the tagged cell types compared to all root transcripts (Figure 3). Annotation clusters related to transcriptional regulation had the highest scores with particularly strong enrichment of transcription factors (TFs) that contain ‘ethylene-responsive elements’ (AP/ERF). Another set of enriched annotations is linked to redox biology and iron, which could be linked to ROS signaling in the tagged cell types. Additional significant clusters contain gene functions related to auxin signaling, post-embryonic morphogenesis and oligopeptide transport as would be expected given the role of the tagged tissues in lateral root development, xylem differentiation and long-distance signaling. Enrichment of glucosinolate biosynthesis genes is consistent with a role of the xylem in long-distance transport of glucosinolates (Andersen et al. 2013). Annotations related to photosynthesis were unexpected but could reflect perception of light piped into the root via the xylem (Nimmo 2018).

**Figure 3:**
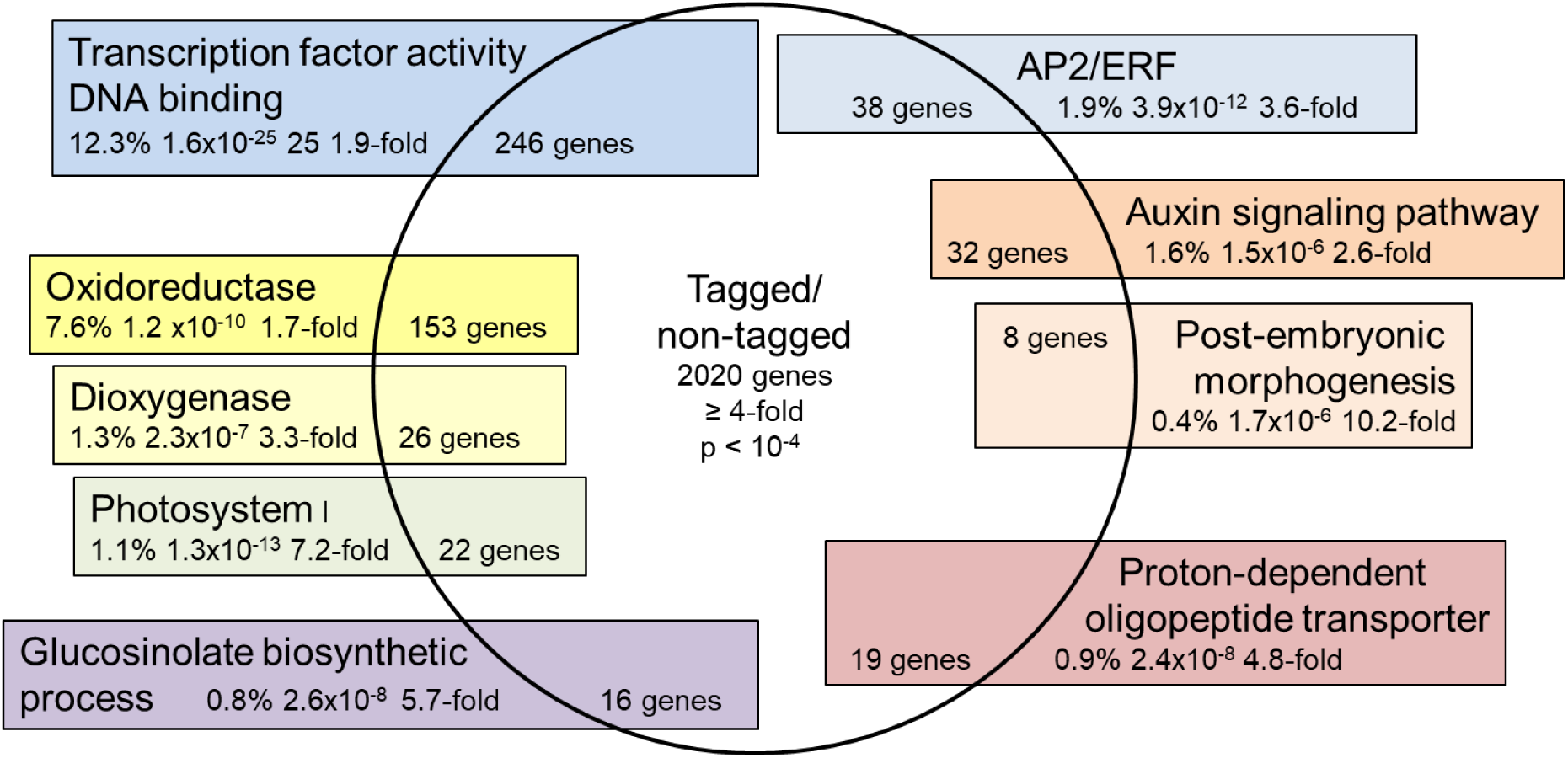
Enrichment of functional gene annotations among genes with preferential expression in the xylem adjacent cell types. Enrichment scores were obtained using DAVID to compare representation of annotation terms in 2020 cell-type specific genes (≥ 4-fold transcript levels in tagged/non-tagged cell samples, p < 10^−4^) with representation in all root expressed genes (25144 genes, see Methods). Names and statistics are only shown for one group in an enrichment cluster.

### H3K27me3 distinguishes HKT1-expressing cell types from whole-root samples

As well as providing a new resource for mining root cell-type specific gene functions, the dataset offered a starting point for investigating causal pathways underpinning and arising from cell-type specific gene expression. To identify genes with differential histone modification patterns in the pHKT1 tagged cell types we performed Isolation of Nuclei Tagged in Specific Cell Types (INTACT; Deal and Henikoff, 2011) and subsequent ChIP-sequencing on roots harvested from 2-weeks old HKT1-INTACT plants. Each of the five replicate samples represented approximately 1600 plants grown in independent batches. Each sample was split and subjected to nuclei isolation either including a procedure to pull down biotinylated nuclei with streptavidin (‘HKT1’ samples) or not (‘Whole Root’ samples). To confirm cell-type specific enrichment of nuclei we spiked the HKT1 samples with root tissue from *A. thaliana* Ler plants (Moreno-Romero et al., 2016; Moreno-Romero et al., 2017). The ratio of Col-0/Ler DNA increased after pull-down, indicating 85-98% purity of pHKT1-NTF nuclei (Supplemental Figure 1). Microscopy of the nuclear preparations further confirmed isolation of GFP-labelled nuclei from the HKT1-expressing cell types (Supplemental Figure 1). Each nuclear sample was further sub-divided and subjected to ChIP with antibodies for H3, H3K4me3 or H3K27me3, followed by Illumina sequencing. The genome-wide profiles of H3, H3K4me3 and H3K27me3 profiles in HKT1 and Whole Root were similar to those previously reported for Arabidopsis roots (Sani et al., 2013).

For quantitative comparison we determined normalised cumulative ChIP-sequence reads over the coding and immediate (200 bp) upstream sequence of each gene (see Methods). For H3K27me3, PCA based on these values separated HKT1 and Whole Root samples along PC2 explaining 14% of the variance (Figure 4). One HKT1/Whole Root sample pair had produced lower total read numbers than the other samples, and this technical difference dominated PC1. PCA without these samples further unmasked the cell-type effect, now separating HKT1 and Whole Root samples along PC1 explaining 72% of the variance (inset in Figure 4). We conclude that H3K27me3 coverage of individual genes is a distinguishing feature of the root cell-types. By contrast, PCA based on H3 or H3K4me3 levels did not separate HKT1 and Whole Root samples (Supplemental Figure 2).

**Figure 4.**
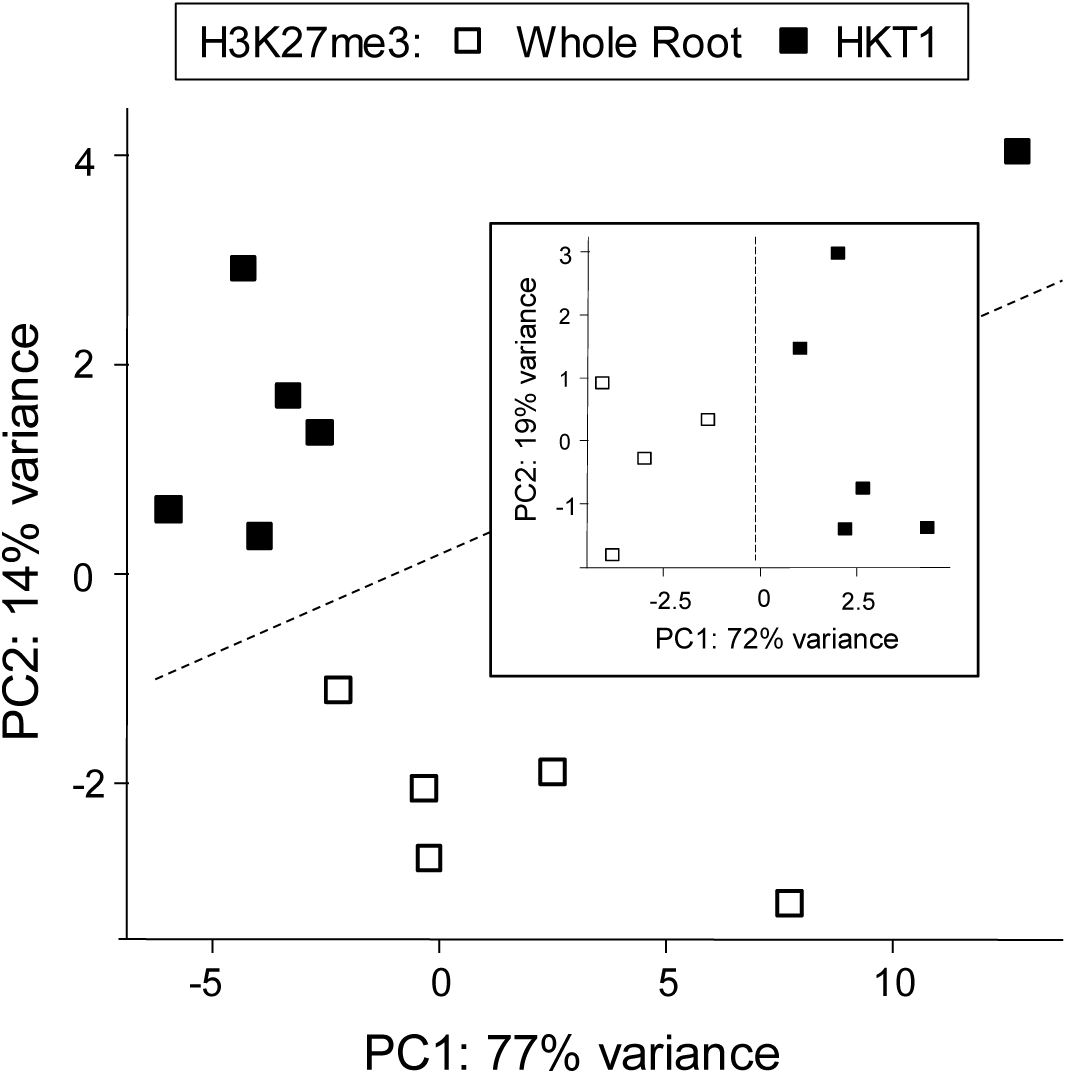
Principal component analysis of genome-wide H3K27me3 levels distinguishes between root cell types. Biotin-labelled nuclei (‘HKT1’, black symbols) were isolated from roots of *A. thaliana pHKT1::NTF pACT2::BirA* lines using pulldown with streptavidin (INTACT). Total nuclear preparations (not subjected to INTACT) from the same root material represent all cell types (‘Whole Root’, open symbols). Replicate samples were obtained from five independently grown plant batches. A sixth sample only subjected to INTACT is also included. The nuclear isolates were subjected to ChIP with antibodies against H3K27me3. The PCA plot shown is based on cumulative H3K27me3 reads over gene bodies (see Methods). PC2 separates HKT1 samples from Whole Root samples explaining 14% of variation. Exclusion of one replicate pair with low total read number (insert) enhanced the distinction between cell types (PC1 explaining 72% of variation).

### Genes with cell-type specific differences in H3K27me3 coverage have distinct functions

The distinction of the cell-types by PCA was based on small but consistent differences in gene coverage with H3K27me3. To identify the differentially marked genes we used a sensitive ranking-based method (Rank Products; Breitling et al., 2004). 168 genes showed significantly lower H3K27me3 and 263 genes showed significantly higher H3K27me3 coverage in HKT1 than in Whole Root samples (FDR < 0.05). Very few genes showed significant differences in H3 levels (1 down and 19 up in HKT1 versus Whole Root,), and none of them showed also differential H3K27me3. Therefore, the identified differences in H3K27me3 cannot be explained with a gain or loss of nucleosomes. Similarly, very few genes had significant differences in H3K4me3 (6 up and 4 down in HKT1 versus Whole Root), and only two of them differed also in H3K27me3 (H3K4me3 up and H3K27me3 down). In summary, H3K27me3 was the most consistent difference between the cell types but only affected a small percentage of the genome.

Based on annotation information for all differentially H3K27me3-marked genes enrichment analysis with DAVID (Huang et al., 2009) revealed distinct functions of genes with cell-type specific differences in H3K27me3 (Figure 5).

**Figure 5:**
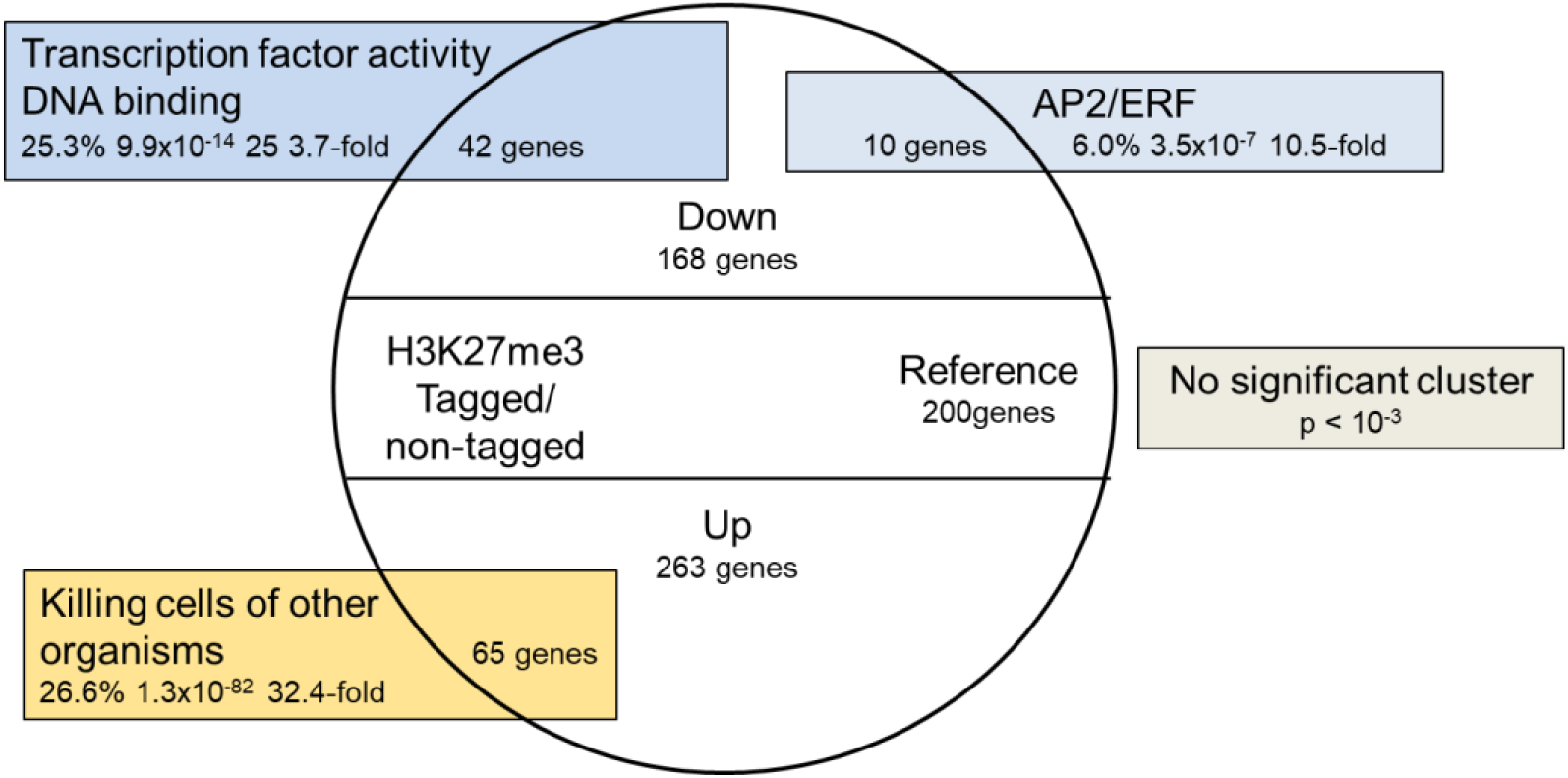
Enrichment of functional gene annotations among genes with lower or higher H3k27me3 levels in xylem-adjacent cell types. Enrichment scores were obtained using DAVID. Gene sets tested contained genes with either significantly lower (Down) or significantly higher (Up) H3K27me3 levels. A similar sized reference set of genes with no changes showed no significant enrichment of functional annotations. Names and statistics are only shown for one group in an enrichment cluster.

Genes with lower H3K27me3 levels in HKT1 compared to Whole Root samples were enriched for transcription factors, listed in Table 1, particularly those containing AP2/ERF domains. For genes with higher H3K27me3 in HKT1 compared to Whole Root samples a significantly enriched cluster contained annotation terms related to antimicrobial, antifungal, and defense functions. A reference set of 200 genes randomly taken from genes with similar H3K27me3 levels in HKT1 and Whole Root samples showed no significant enrichment (p < 10^−3^) of functional annotations.

**Table 1.**
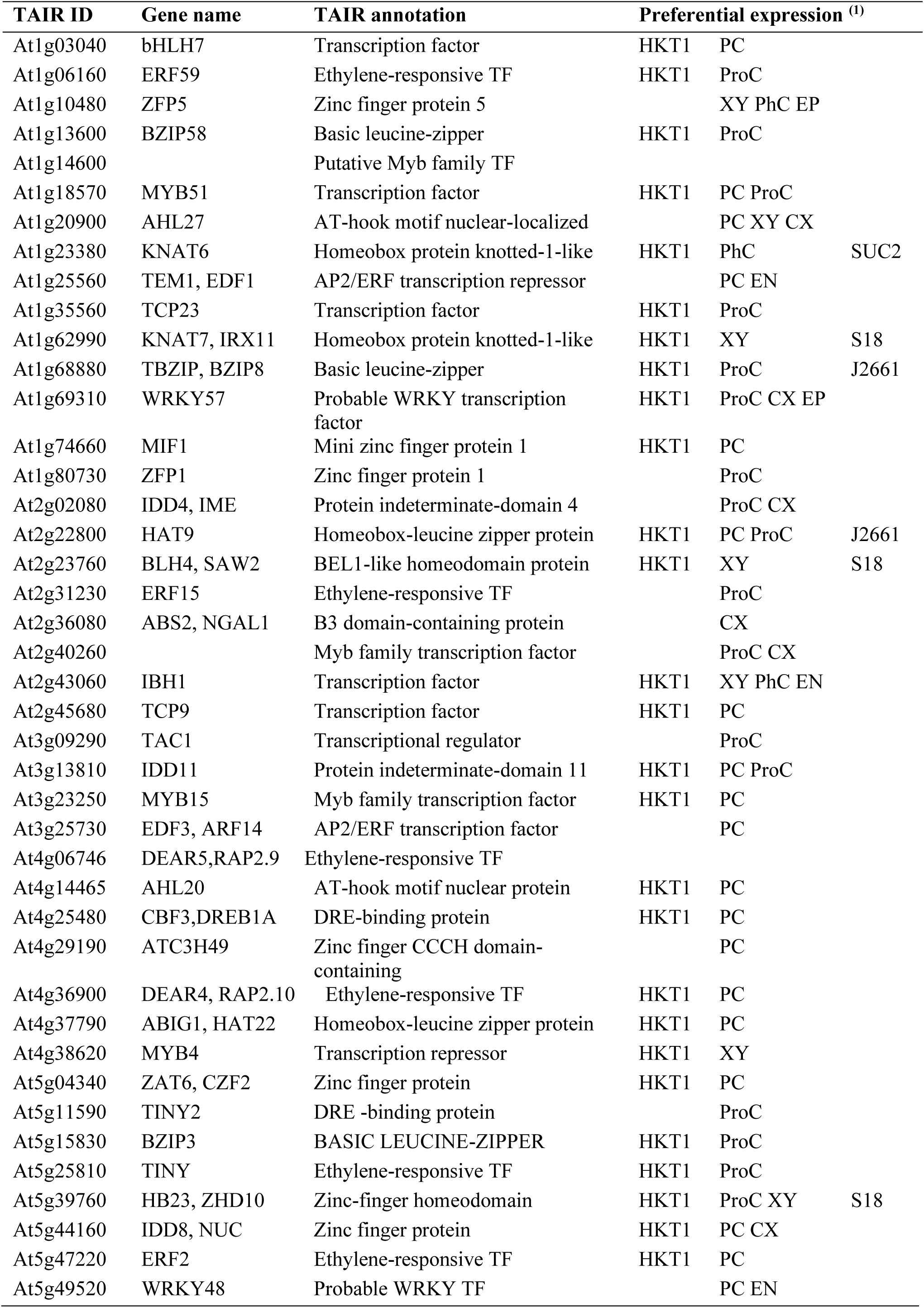
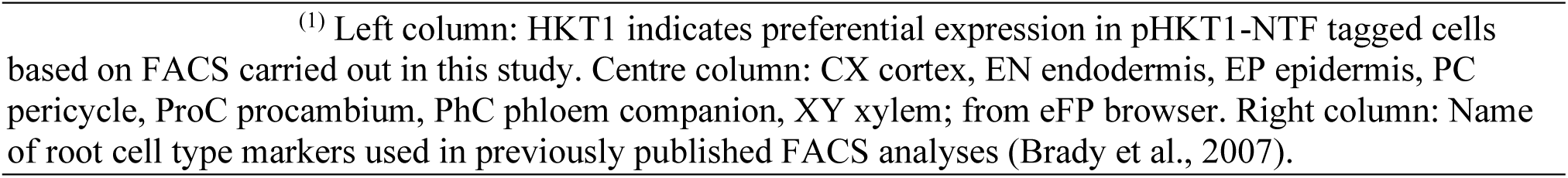
Transcription factors with significantly lower H3K27me3 levels in ‘HKT1’ cell types.

### Differentially H3K27me3 marked genes have tissue-specific expression patterns

H327me3 is a repressive mark, and we therefore asked whether genes with lower H3K27me3 levels had higher transcript levels in the tagged cell types. We consulted both previously published data (Birnbaum et al., 2003; Brady et al., 2007) and our pHKT1-NTF based FACS dataset. Gene lists based on previously employed root cell-markers yielded very few hits, emphasizing the fact that these studies did not include specific markers for the root tissues where pHKT1 is active. Analysis of computationally inferred expression profiles available in the eFP browser (Winter et al., 2007) indicated preferential expression in the xylem-adjacent cell types for the majority of the 168 H3K27me3-depleted genes. For example, 30 of them are displayed in the eFP browser with exclusive expression in the pericycle and 54 with exclusive expression in the ‘procambium’, the meristematic tissue that produces the xylem parenchyma. A reference subset of 200 genes randomly picked from genes with equal H3K27me3 levels in HKT1 and Whole Root samples showed no such tissue bias in the eFP browser.

Comparison of the 168 H3K27me3-depleted genes with our own FACS results identified a set of 86 genes that had both significantly lower H3K27me3 and significantly higher transcript levels in the tagged cell types than in the rest of the root (Figure 6). The 51% overlap (86/168 genes) was significantly higher (p=1.5×10^−44^) than the 8% (2020/25144 genes) expected for any random subset of genes. Indeed, overlap of the 2020 preferentially expressed genes with the 200 genes in the reference set contained only 16 genes (8%) and overlap with the list of 263 genes with higher H3K27me3 contained only 4 genes (1.5%). Only one of the 168 H3K27me3-depleted genes showed a lower transcript level in tagged cells compared to non-tagged cells. 27 (64%) of the 42 H3K27me3-depleted transcription factors (Table 1) were preferentially expressed in the tagged cell types, representing 31% of the 86 shared genes, which is significantly higher than the percentage of TFs in the whole genome (8%), or among the 2020 cell-type specific genes (10%) with p-values of 5×10^−20^ and 5×10^−18^ respectively.

**Figure 6.**
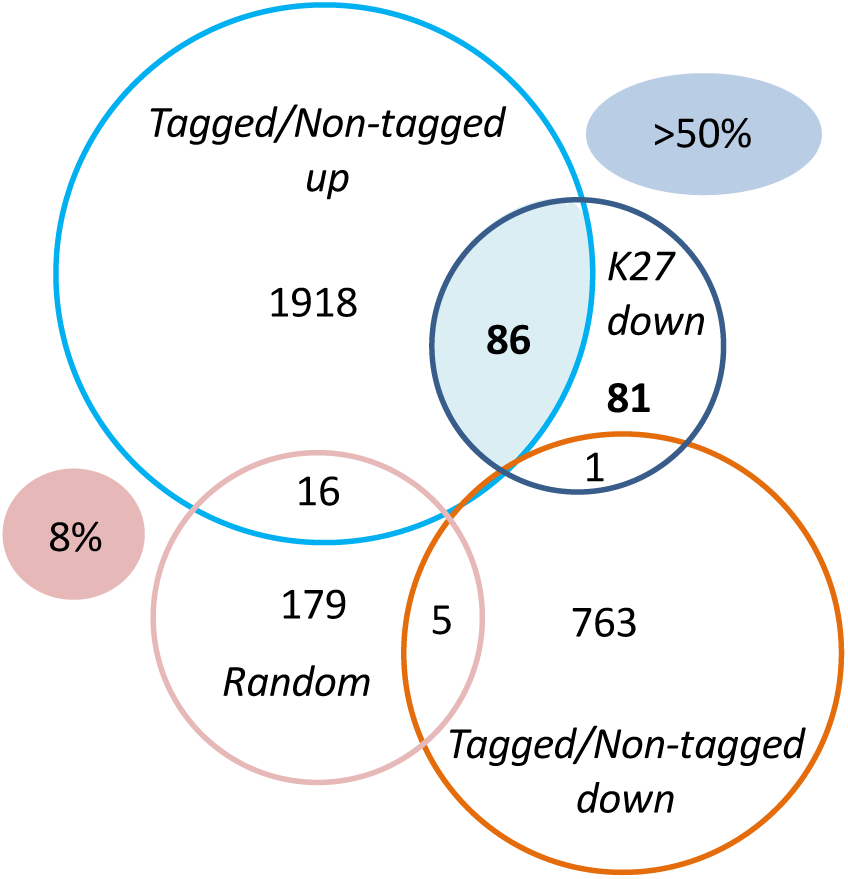
Overlap between genes with lower H3K27me3 levels and genes with preferential expression in the tagged cell types. Venn diagrams are based on lists of genes identified with FACS or INTACT. The genes with higher transcript level in tagged versus non-tagged cells (2020 genes, light blue circle) represent approximately 8% of the total transcriptome. The overlap between a random set of 200 genes with no change in H3K27me3 (pink circle) and the cell-type specific expressed genes (light blue circle) is 8% (16 genes). By contrast, 51% of genes with a cell-type specific decrease in H3K27me3 level (86 of 168 genes, dark blue circle) are preferentially expressed in the tagged cell types. Very little overlap (1 gene) was found with genes showing lower expression in tagged over non-tagged cell types (769 genes, brown circle).

Overall, the analysis revealed a statistical association between cell-type specific loss of H3K27me3 and increased gene expression, and a strong bias of the cell-type specific epigenetic de-repression for transcription factors. The 27 TFs (labelled ‘HKT1’ in Table 1) are therefore good candidates for setting up local transcriptional networks. 14 of them were significantly upregulated by a 24-h salt treatment of the roots (Figure 7).

**Figure 7.**
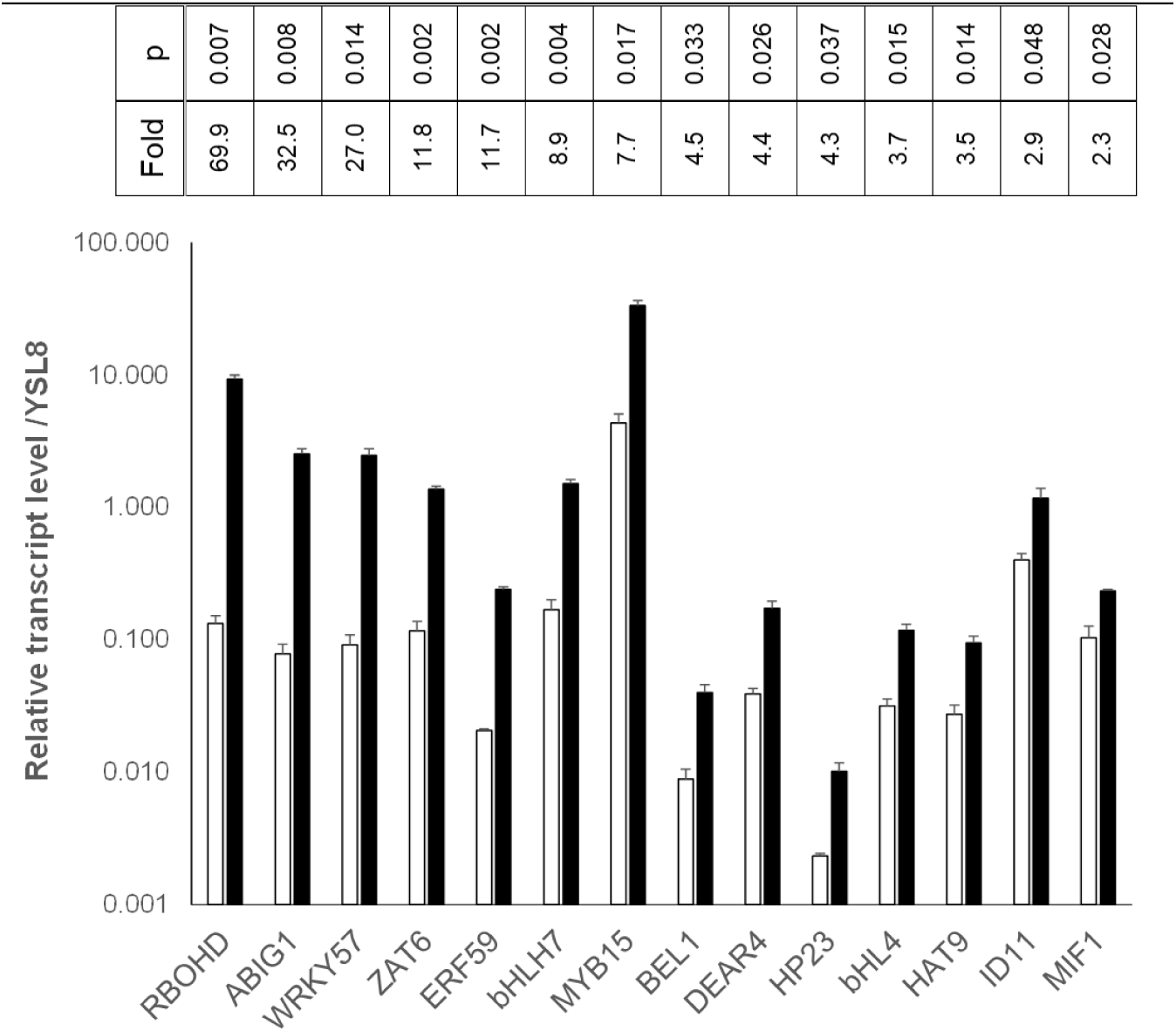
Cell-type specifically de-repressed transcription factors are regulated by salt. Transcript levels of the genes listed in Table 3 levels were determined by RT-qPCR. 14 of them showed significant induction by salt. pHKT1-NTF*:p*ACT2*-BirA* plants were grown in hydroponics and treated with 150 mM NaCl for 24 hours (black bars), or not (open bars). Each biological replicate represents RNA harvested from 10 roots, grown in independent batches. Bars are means of transcript levels across 3 biological replicates, error bars are SE. Fold-changes and p-values (paired student’s t test) are listed above bars. Within each biological replicate, transcript levels were normalised to reference gene YSL8 (At1g48370) in two technical replicates.

### The cell-type specific transcription factor ZAT6 is de-methylated by REF6

H3K27me3 levels are determined by the relative rate of methylation and demethylation through methyltransferases and demethylases, respectively. REF6 (At3g48430) encodes one of the H3K27me3-specific demethylases in *A. thaliana* (Lu et al., 2011; Yan et al., 2018). Comparison of the cell-type specifically de-repressed transcription factors (Table 1) with a published list of demethylated genes in whole seedlings of *ref6* mutants (Lu et al., 2011) highlighted ZAT6 (At5g04340) as a potential REF6-target. To test REF6-dependence of ZAT6 we performed anti-H3K27me3 ChIP-Seq on mature roots of wildtype and *ref6* plants, as well as ChIP-qPCR and RT-qPCR. The genome-wide ChIP patterns in Figure 8 show a distinct H3K27me3 island covering the ZAT6 coding sequence. Coverage is decreased in the HKT1 samples compared to Whole Root samples (A) and increased in *ref6* mutant compared to wildtype (B). ChIP-qPCR and RT-qPCR experiments presented in Figure 9 showed significantly higher ZAT6 H3K27me3 levels (A) and lower ZAT6 transcript levels (B) in two independent *ref6* mutant compared to wildtype (B), indicating that ZAT6 expression in roots depends on REF6-mediated H3K27me3 demethylation.

**Figure 8.**
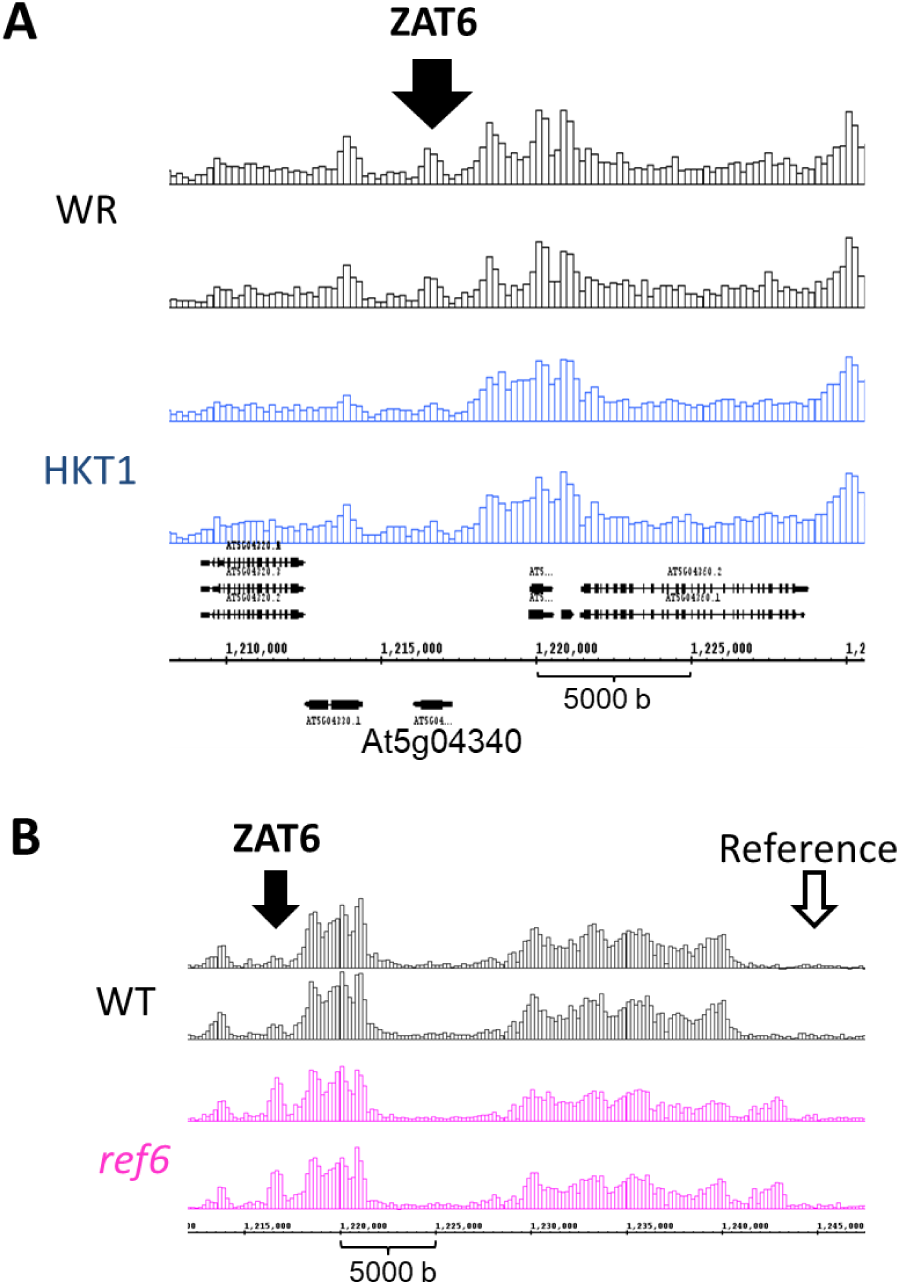
H3K27me3 coverage of ZAT6 is decreased in pHKT1-NTF tagged cell types and increased in *ref6* mutants. **B:** H3K27me3 coverage over ZAT6 in roots of Whole Root (WR) and HKT1 INTACT samples. For each genotype two independent replicates are shown. Positions of TAIR gene models are shown under the ChIP-seq reads. **B:** H3K27me3 profiles in roots of *A. thaliana* wildtype and *ref6-3* plants obtained from ChIP-seq reads in 200 bp-windows. For each genotype two independent replicates are shown. Positions of ZAT6 and a reference region used for ChIP-qPCR are indicated with black and white arrows respectively.

**Figure 9.**
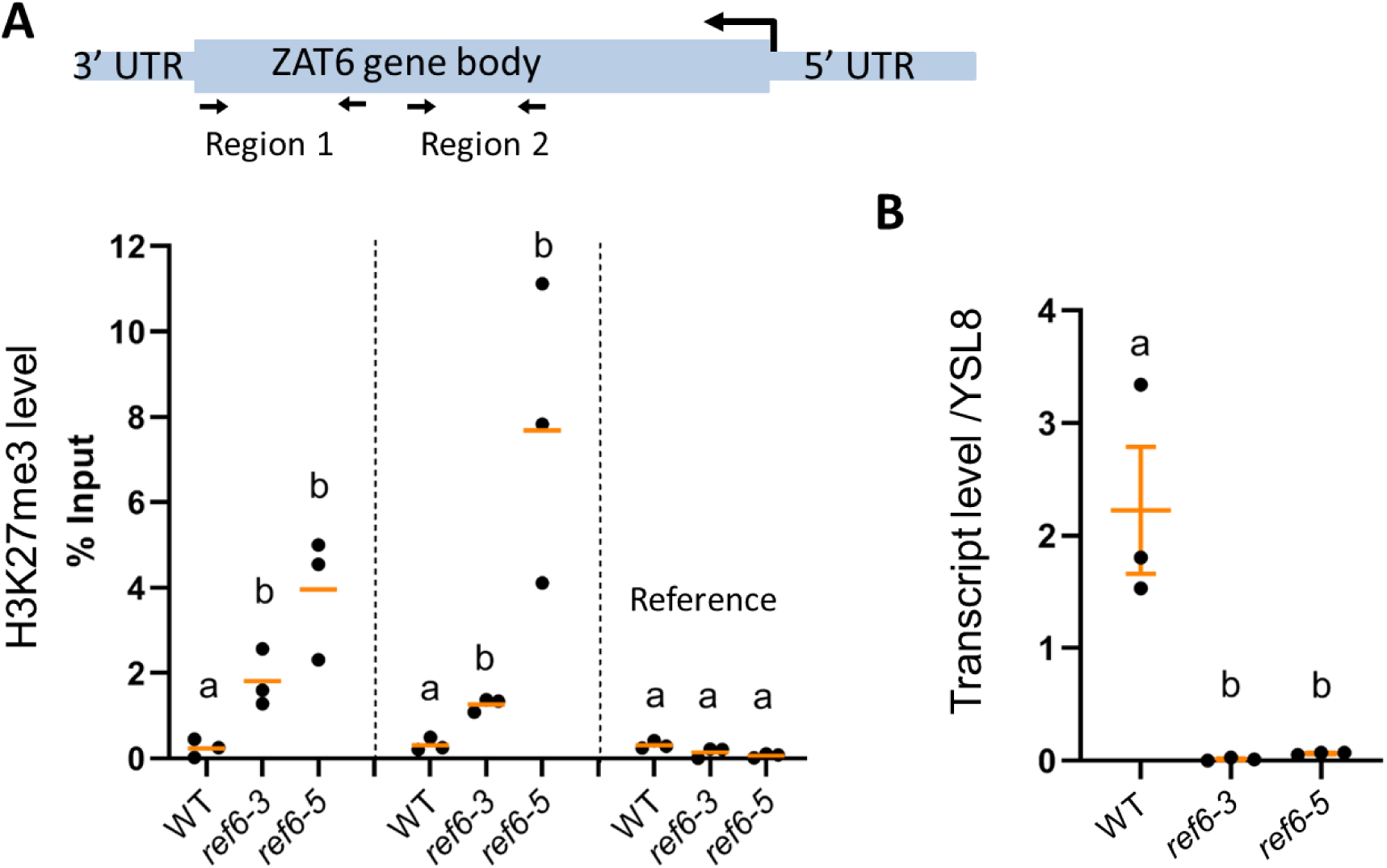
H3K27me3 and transcript levels of ZAT6 depend on the histone demethylase REF6. **A:** H3K27me3 levels (relative to Input) in regions 1 and 2 of ZAT6, and in the reference region, in roots of wildtype, *ref6-3* and *ref6-5* mutant lines as determined by anti-H3K27me3 ChIP-qPCR. Dots represent the results from three independently grown plant batches. Means are shown as orange lines. Different letters indicate significant differences (paired Student’s t-test, p<0.05). **B:** ZAT6 transcript levels in roots of wildtype, *ref6-3* and *ref6-5* mutant lines as determined by RT-qPCR. Data are relative to the constitutively expressed YSL8. Dots represent the results from three independently grown plant batches. Means are shown as orange lines. Different letters indicate significant differences between genotypes (paired t-test, p<0.05).

### Many cell-type specific transcripts are downstream of ZAT6

To test whether ZAT6 could establish a cell-type specific downstream network we analysed root transcriptome of the two *zat6* mutant lines by RNA-sequencing. PCA clearly distinguished between mutant and wildtype samples based on normalized transcript levels (Figure 10). Wildtype and mutant samples were separated by PC1 explaining 78% of variance. The replicates of each mutant line also grouped closely together, allowing distinction between the two lines primarily through PC2 (explaining 5% of variance). 349 genes showed a significant decrease in transcript level (> 2-fold, p < 10^−3^) in both *zat6* mutant lines compared to wildtype. Comparison with the FACS dataset revealed that most of the ZAT6-dependent genes were preferentially expressed in the tagged cell types (Figure 11).

**Figure 10.**
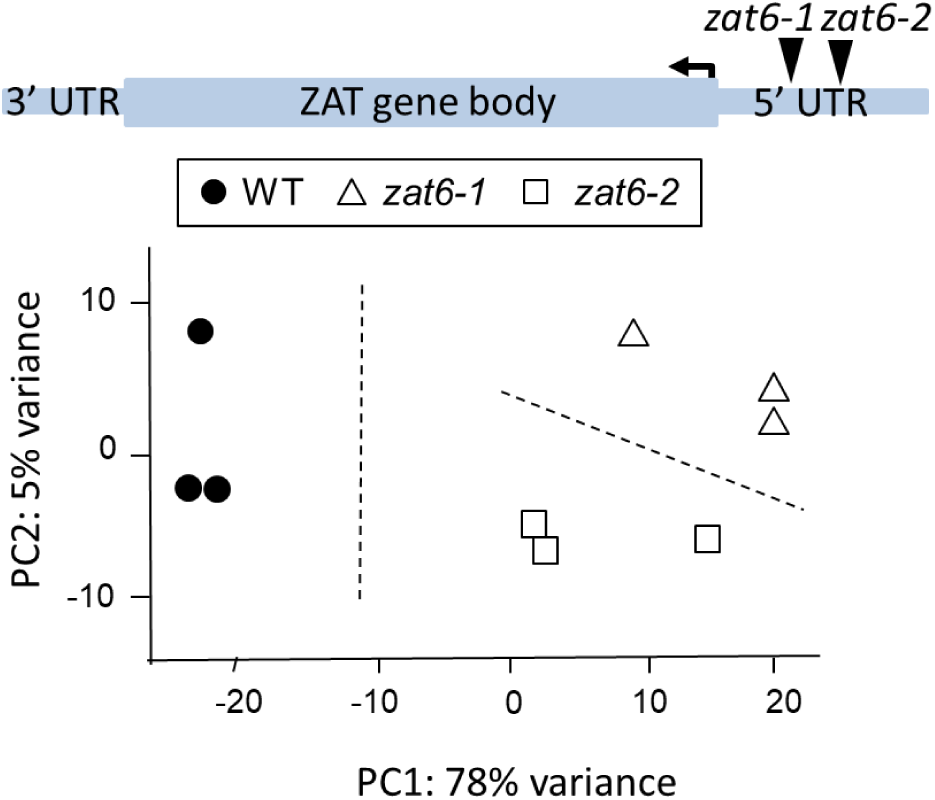
ZAT6 mutant transcriptomes are significantly different from wildtype. PCA based on genome wide transcript levels in roots of *A. thaliana* wildtype (black circle) and two *zat6* mutants (open symbols). Replicate samples were obtained from three independently grown plant batches for each genotype. The ZAT6 gene model on the top shows the position of the T-DNA insertion for the two independent *zat6* mutant lines.

**Figure 11.**
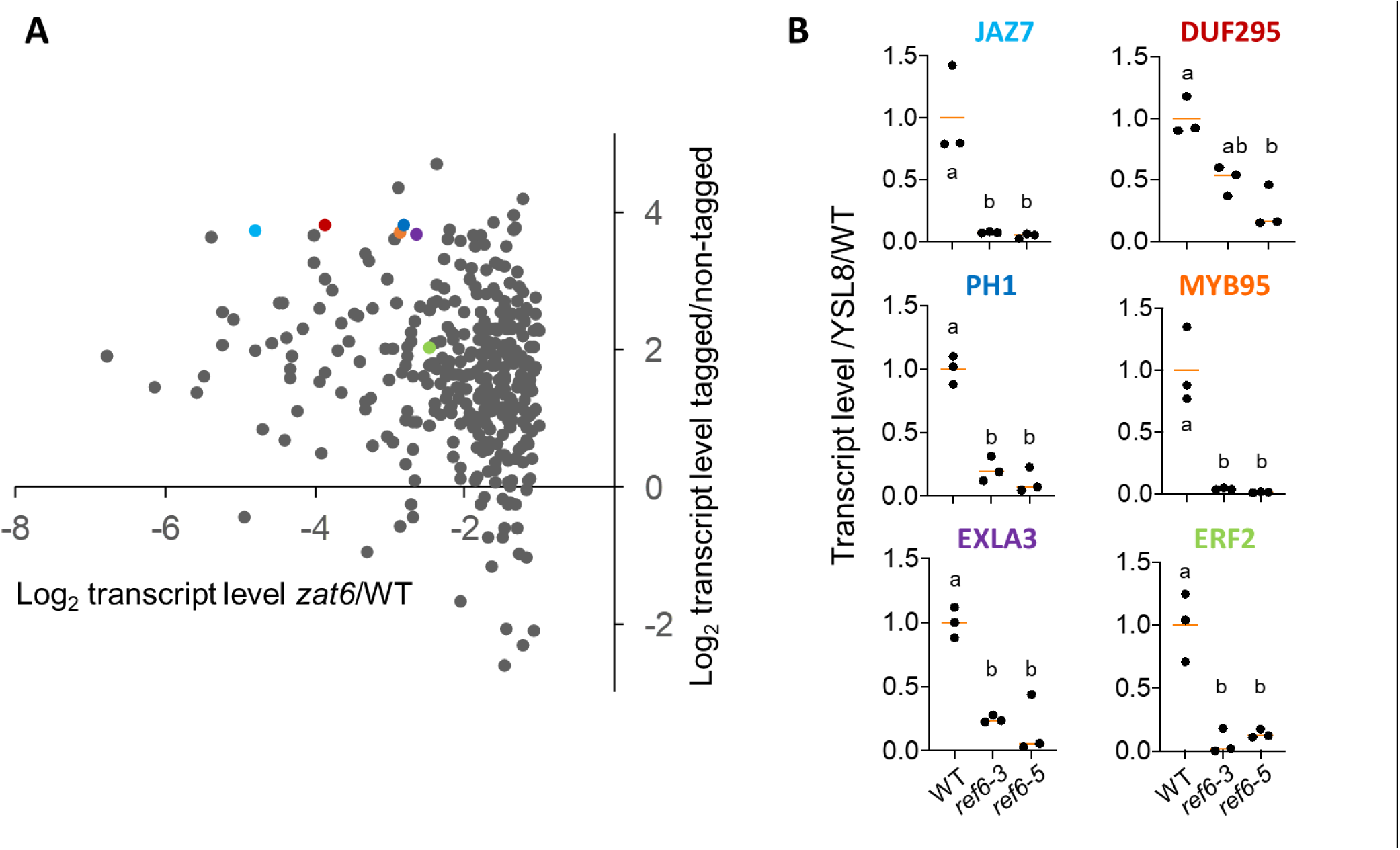
The ZAT6-regulome is cell-type biased and REF6-dependent. **A:** Ratio of transcript levels in the pHKT1-NTF tagged cells (T) versus non-tagged root cells (NT) in wildtype plants (as determined by FACS-RNA-seq, Supplementary Dataset 1) is plotted against the average ratio of transcript levels in *zat6* lines versus wildtype (log_2_ of mean transcript level in *zat6-1* and *zat6-2* divided by mean transcript level in wildtype, WT). Values are shown for 349 genes with significantly decreased levels in both *zat6* lines (>2-fold, p<10^−3^; Supplementary Dataset 10). Couloured dots indicate the genes analysed by qPCR (B) **B:** Transcript levels of ZAT6-dependent genes in roots of wildtype (WT), *ref6-3* and *ref6-5* mutant lines as determined by RT-qPCR. |Gene name colour indicates the respective data points in (A). Transcript levels are relative to YSL8 and normalised to the average WT value. Dots represent the results from three independently grown plant batches. Means are shown as orange lines. Different letters indicate significant differences between genotypes (paired Student’s t-test, p<0.05).

Plotting for each gene the mean relative transcript level in *zat6*/WT against the relative transcript level in tagged/non-tagged cell types shows a clear bias towards cell-type specific expression among the ZAT6-dependent genes (Figure 11 A). 105 (30%) of the ZAT6-dependent genes met the stringent cut-off criteria for preferential cell-type expression applied to the FACS dataset (at least 4-fold, p < 10^−4^), significantly more than the 8% expected by random (4×10^−33^). We conclude that cell-type specific de-repression of ZAT6 proliferates gene expression in a largely tissue-delimited fashion thus establishing one branch of the local transcriptional network.

To verify that H3K27me3-demethylation of ZAT6 is critical for the expression of other genes in this branch, we measured transcript levels of ZAT6-dependent cell-type specific genes (coloured in Figure 11A) in *ref6* mutants using RT-qPCR. All of the genes tested lost expression in the *ref6* mutant lines (Figure 11B). According to previous published data (Lu et al., 2011) and our own *ref6* ChIP-Seq experiment, these genes are not direct REF6-targets. Hence their REF6-dependence suggests indirect regulation via H3K27me3 de-methylation of ZAT6.

Functional annotation enrichment analysis of ZAT6-dependent genes with preferential expression in the tagged root cell types identified a significant annotation cluster related to jasmonic acid, wounding and defense signaling. The core gene sets underpinning this cluster contained several JAZ proteins and oxylipin biosynthesis enzymes (Table 2). Transcription factors, particularly those with AP2/ERF elements, were also again over-represented.

**Table 2.**
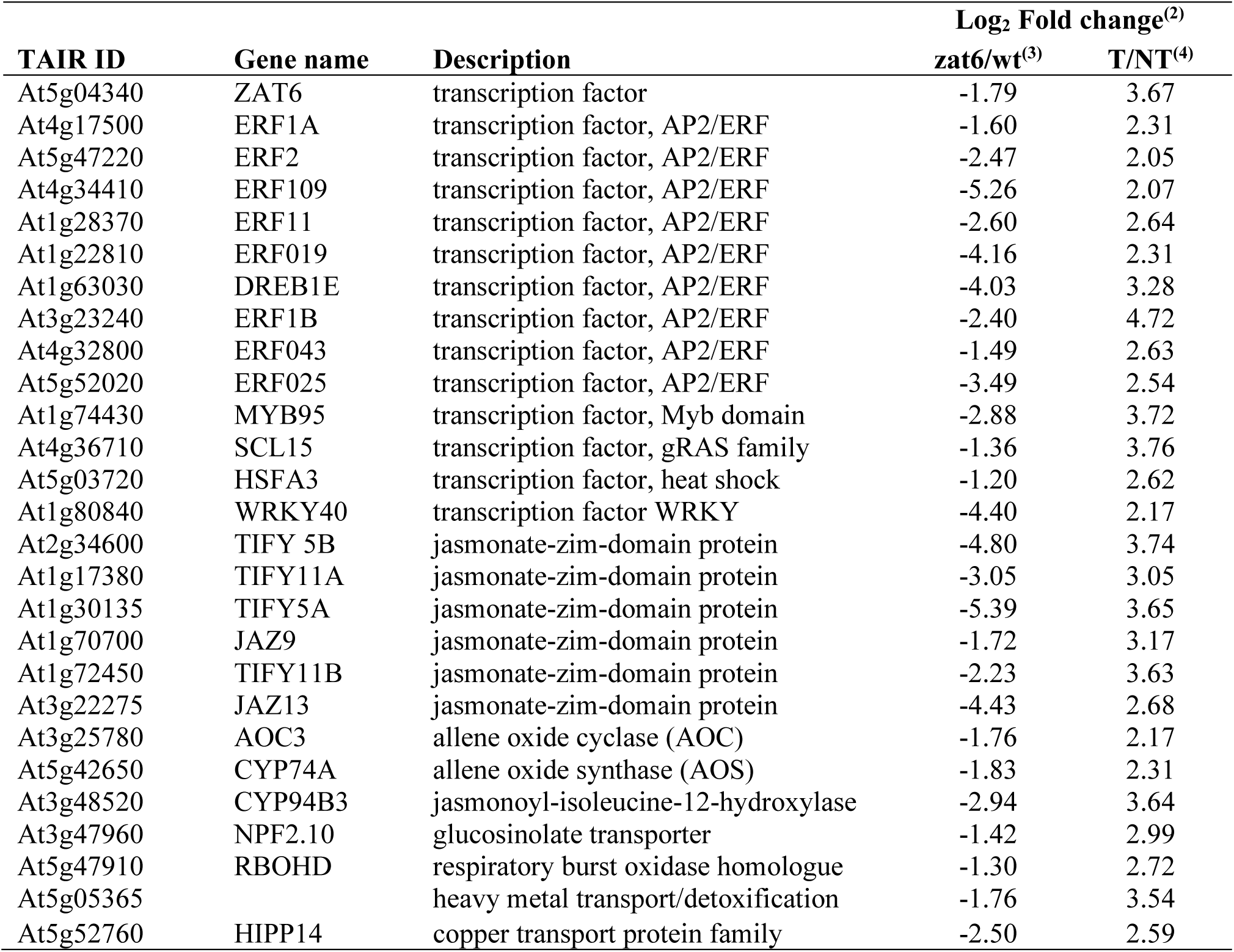
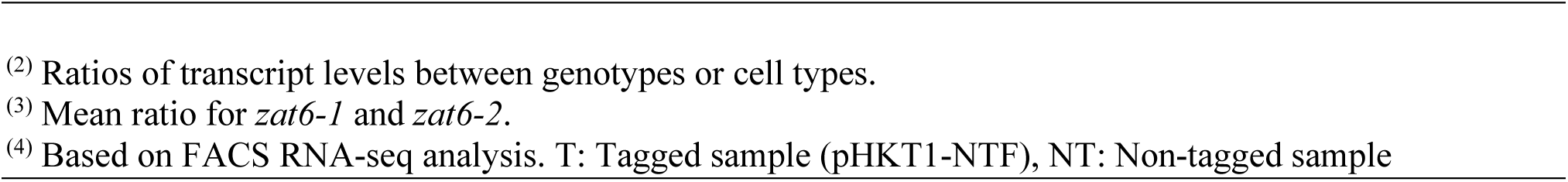
Examples of ZAT6-dependent genes preferentially expressed in pHKT1-NTF tagged cell types.

| TAIR ID | Gene name | Description | Log<sub>2</sub> Fold change<sup>(2)</sup> |  |
| --- | --- | --- | --- | --- |
|  |  |  | *zat6*/wt<sup>(3)</sup> | T/NT<sup>(4)</sup> |
| At5g04340 | ZAT6 | transcription factor | -1.79 | 3.67 |
| At4g17500 | ERF1A | transcription factor, AP2/ERF | -1.60 | 2.31 |
| At5g47220 | ERF2 | transcription factor, AP2/ERF | -2.47 | 2.05 |
| At4g34410 | ERF109 | transcription factor, AP2/ERF | -5.26 | 2.07 |
| At1g28370 | ERF11 | transcription factor, AP2/ERF | -2.60 | 2.64 |
| At1g22810 | ERF019 | transcription factor, AP2/ERF | -4.16 | 2.31 |
| At1g63030 | DREB1E | transcription factor, AP2/ERF | -4.03 | 3.28 |
| At3g23240 | ERF1B | transcription factor, AP2/ERF | -2.40 | 4.72 |
| At4g32800 | ERF043 | transcription factor, AP2/ERF | -1.49 | 2.63 |
| At5g52020 | ERF025 | transcription factor, AP2/ERF | -3.49 | 2.54 |
| At1g74430 | MYB95 | transcription factor, Myb domain | -2.88 | 3.72 |
| At4g36710 | SCL15 | transcription factor, gRAS family | -1.36 | 3.76 |
| At5g03720 | HSFA3 | transcription factor, heat shock | -1.20 | 2.62 |
| At1g80840 | WRKY40 | transcription factor WRKY | -4.40 | 2.17 |
| At2g34600 | TIFY 5B | jasmonate-zim-domain protein | -4.80 | 3.74 |
| At1g17380 | TIFY11A | jasmonate-zim-domain protein | -3.05 | 3.05 |
| At1g30135 | TIFY5A | jasmonate-zim-domain protein | -5.39 | 3.65 |
| At1g70700 | JAZ9 | jasmonate-zim-domain protein | -1.72 | 3.17 |
| At1g72450 | TIFY11B | jasmonate-zim-domain protein | -2.23 | 3.63 |
| At3g22275 | JAZ13 | jasmonate-zim-domain protein | -4.43 | 2.68 |
| At3g25780 | AOC3 | allene oxide cyclase (AOC) | -1.76 | 2.17 |
| At5g42650 | CYP74A | allene oxide synthase (AOS) | -1.83 | 2.31 |
| At3g48520 | CYP94B3 | jasmonoyl-isoleucine-12-hydroxylase | -2.94 | 3.64 |
| At3g47960 | NPF2.10 | glucosinolate transporter | -1.42 | 2.99 |
| At5g47910 | RBOHD | respiratory burst oxidase homologue | -1.30 | 2.72 |
| At5g05365 |  | heavy metal transport/detoxification | -1.76 | 3.54 |
| At5g52760 | HIPP14 | copper transport protein family | -2.50 | 2.59 |

### ZAT6 restricts sodium accumulatio in the shoots

Spatial co-expression of ZAT6 with HKT1, up-regulation by salt, and over-representation of ERFs in the downstream network prompted us to investigate a possible role of ZAT6 in regulating long-distance transport of Na. Figure 12 shows Na and K concentrations in leaves of wildtype and *zat6* plants after 3 days exposure to 100 mM NaCl. As expected, all salt-treated plants accumulated Na in the shoots but Na concentrations were more variable and on average significantly higher in *zat6* mutant plants than in wildtype plants. No differences were found for K. These results highlight a function of ZAT6 in regulating root-to-shoot Na allocation under salt stress.

**Figure 12.**
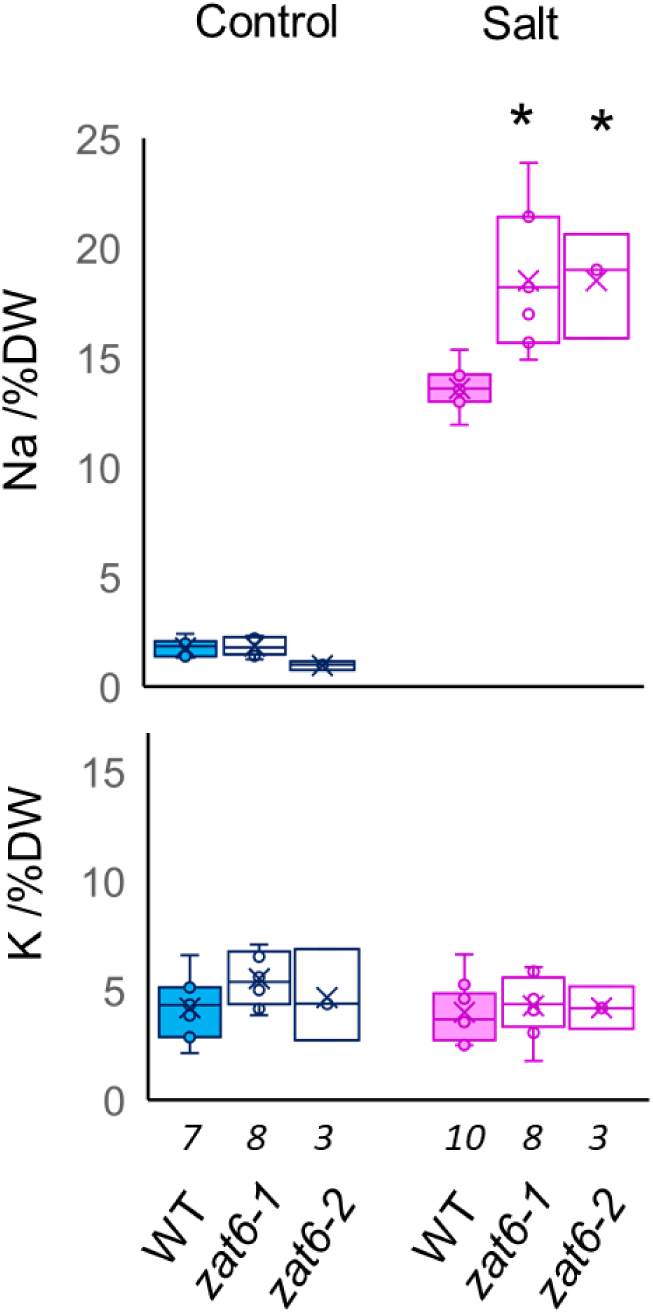
ZAT6 limits sodium accumulation in the shoots. Shoot concentrations (%DW) of Na (top) and K (bottom) in wildtype (filled bars) and two independent *zat6* lines (open bars) treated with 100 mM NaCl for 5 days (Salt) or not (Control). Box blots show means and standard deviation of values from individual, independently treated, plants (n given under genotype). Asterisks indicate significant difference between salt-treated *zat6* and salt-treated wildtype (p < 0.01). All genotypes had significantly higher Na shoot concentrations in salt compared to control (p < 0.001).

## DISCUSSION

### HKT1 co-transcriptome and histone methylation patterns in xylem-adjacent cell types

The focus of this study was on cell types that control long-distance transport and communication between the roots and the shoots, which is an essential prerequisite for adjusting shoot growth and development to the presence of water, nutrients and stress factors in the soil environment. Previous studies used root cell-type markers that are primarily expressed in the tips where the xylem cells are still alive. However, the content of mature (dead) xylem vessels in the upper part of the root is controlled by the adjacent cell layers. This was nicely shown by studies on the transporter HKT1. HKT1 prevents accumulation of toxic Na^+^ levels in the shoots by retrieving Na^+^ from the transpiration stream (Munns et al., 2012; Munns and Tester, 2008), and its specific expression in the xylem parenchyma and pericycle of mature roots (Sunarpi et al., 2005) is critical for function (Møller et al., 2009). HKT1 is therefore an excellent marker for the cell layers that act as a checkpoint for exchange of substances between the endodermis-controlled symplast of the stele and the apoplastic xylem vessels (Barberon and Geldner, 2014). The pericycle has the additional function to generate lateral roots, which is reflected in root architectural phenotypes of *hkt1* mutants (Julkowska et al., 2017).

The pHKT1:NTF lines generated here showed strong and specific GFP fluorescence in the cell layers surrounding mature xylem vessels (Figure 1). FACS-RNA-Seq analysis identified over 2000 genes with preferential expression in the HKT1-expressing cells. This new dataset represents a foundation for further studies into specific gene functions in these tissues. All raw data have been made publicly available for further data mining and specific gene lists with functional annotations can be requested from the authors by interested parties. To facilitate characterization of cell-type specific regulatory pathways we looked for potential primary regulators that do not require upstream TFs for cell-type specific expression. In particular, we tested whether a cell-type specific loss of H3K27me3 in a few genes could potentially underpin the entire cell-type specific transcriptome. Integration of INTACT ChIP-Seq data with FACS RNA-Seq data identified a set of genes with both lower H3K27me3 and higher expression in the HKT1-tagged cell types, highly enriched for transcription factors (Table 1). Preferential targeting of transcription factors for cell-type specific changes in H3K27me3 was also found in guard-cell lineages (Lee et al., 2019) and emerges as a candidate mechanism for establishing cell-type specific regulatory networks.

### The ZAT6-branch of the HKT1 co-transcriptome

To test whether epigenetically de-repressed TFs could establish larger cell-type specific transcriptional networks we further interrogated upstream and downstream regulation of the transcription factor ZAT6. ZAT6 is preferentially expressed in the tagged cell-types and shows cell-type specific decrease of H3K27me3 (Table 1, Figures 8,9). Analysis of *ref6* and *zat6* mutants showed that that (i) REF6 mediates H3K23me3 demethylation of ZAT6 (Figure 8A, 9A), (ii) REF6 is required for expression of ZAT6 (Figure 9B), (iii) a large number of genes are transcriptionally dependent on ZAT6 (Figure 10), most of which show cell-type specific expression (Fig. 11A), and (iv) REF6-dependence of ZAT6 expression is levied onto downstream gene that are not direct REF6-targets (Figure 11B). These results provide proof-of-concept that cell-type specific epigenetic de-repression of a single transcription factor can establish cell-type specific expression of a large number of genes. Supplemental Figure 3 summarizes this conclusion in a working model.

### What is upstream of ZAT6 de-repression?

But why is H3K27me3 removal from ZAT6 cell-type specific? Our FACS experiments showed that REF6 itself is not preferentially expressed in the tagged cell-types. ZAT6 is a candidate target of H2A ubiquitination, which can recruit REF6 (Kralemann et al., 2020), Table S5 therein), but neither Ub13 nor Ub14 showed specific expression in the tagged cell types. Alternatively, ZAT6 could be protected against PRC2-mediated H3K27 methylation in a cell-type specific manner. Differential expression patterns of several PRC2 components in the root have been reported (de Lucas et al., 2016), but apart from slightly lower transcript levels of CLF (At2g23380) we did not find specific repression of PRC2 components in the tagged cell-types. Whether the enzymes are cell-type specifically recruited, activated or effective (Wang et al., 2020) remains to be explored.

### Downstream ‘proliferation’ of cell-type specific de-repression of ZAT6

Assessing cell-type specificity of ZAT6 downstream genes through RNA-Seq of *zat6* mutants allowed us to quantitatively interrogate the conservation of spatial expression within transcriptional pathways. We found that at least a third of the ZAT6-dependent transcriptome maintain strong cell-type preference. Preferential expression in the tagged cell types is increasingly lost among genes that are weakly dependent on ZAT6 (Figure 11A), indicating that the downstream cascade eventually expands beyond the cell type. Dilution of cell-type specificity is expected if parallel transcriptional pathways in other cell types target the same genes, or if positional effects or transcript movement come into play. The ZAT6-downsetram transcriptome therefore offers new opportunities to investigate how different spatial cues interact on individual genes.

### ZAT6-dependent gene functions in the xylem adjacent cell layers

Previous studies have reported roles of ZAT6 in chilling, osmotic stress, salt, cadmium, phosphate and pathogen responses as well as flowering (Devaiah et al., 2007; Shi et al., 2014; Chen et al., 2016; Liu et al., 2013). However, this evidence was derived from analysing whole seedlings and shoots of *zat6* mutants, or from ectopic 35S-driven over-expression of ZAT6, and is therefore not informative for cell-type specific function in roots. Expression of the glutathione biosynthesis gene GSH1 (At4g23100), which was identified as a direct target of ZAT6 in heterologous systems and linked to Cd-tolerance, was not ZAT6-dependent in roots. However, two ZAT6-dependent heavy metal-chelating/transport proteins are specifically expressed in the xylem-adjacent cells (AT5G05365, AT5G52760) and should be tested for a role in long-distance transport.

ZAT6 expression has been reported to be up-regulated by biotic and abiotic stress (Devaiah et al., 2007; Shi et al., 2014; Chen et al., 2016; Liu et al., 2013) and we measured up-regulation in the roots by salt (Figure 7). Furthermore, salt-treated *zat6* mutant plants hyperaccumulated Na^+^ in the shoots compared to wildtype (Figure 12). This phenotype mimics that of mutants for ion transporters, such as HKT1 and SOS1 (Shi et al., 2002, Møller et al., 2009) that remove Na^+^ from the root xylem. The ZAT6-dependent transcriptome did not include any known Na^+^ transporter genes, but it contained candidates for their posttranslational regulation. Notably, the type-2C protein phosphatase PP2C49 (At3g62260), previously shown to regulate HKT1 (Chu et al. 2020), was preferentially expressed in the tagged cell types and lost expression in the *zat6* mutants. Our study provides a limited set of spatially co-expressed genes for systematic interrogation of regulatory pathways through reverse genetics.

The functional enrichment analyses of cell-type specific genes that showed both H3K27me3 de-methylation and ZAT6-dependency also pinpointed hormonal signals. While not all AP/ERF TFs are bona fide targets of ethylene, their over-representation ties in with reports on radial ethylene signals controlling xylem content and long-distance transport of Na^+^ and K^+^ (Jiang et al., 2013). A second significantly enriched annotation cluster pointed to ZAT6 mediating cell-type specific jasmonate responses (Table 2) and supports a role of xylem-adjacent cells in communicating wounding and disease in the root to the shoot (Biere and Goverse, 2016).

In summary, our study provides proof-of-concept for establishment of a cell-type specific transcriptional network from an epigenetically regulated TF. Alongside ZAT6, 26 TFs with both cell-type specific expression and cell-type specific H3K27me3-demethylation were identified (Table 1). Assuming a similar-sized ‘followership’ as for ZAT6, they could generate cell-type specific transcriptome of over 2000 genes thereby offering a handle to systematically delineate regulatory modules in root xylem-surrounding cell types from the ‘bottom up’. Understanding the regulatory networks in a strategic location at the root/shoot interface is essential for improving water and nutrient allocation in plants as well as resilience against soil-borne diseases and toxins.

## METHODS

### Generation of the HKT1 INTACT line

The promoter sequence of HKT1 (At4g10310), spanning from -837 bp to + 6 bp relative to TSS (Mäser et al., 2002) was amplified by PCR from *Arabidopsis thaliana* genomic DNA, sub-cloned into the pGEM T-Easy Vector (Promega), amplified with primers *att*B1 5’-TTACTCCATGTGTCAATACCAAAA-3’ and *att*B2 5’-GTCCATTTTAGTTCTCGAGTCGG-3 and introduced into the pDONR207 Gateway donor vector (Invitrogen). The *NTF-GFP* fragment was amplified by PCR from the published construct (Deal and Henikoff, 2010) using the forward primer 5’-atatctagaATGAATCATTCAGCGAAAACCACAC-3’ and reverse primer 5’-atcgagctcTCAAGATCCACCAGTATCCTC-3’, containing restriction sites for the enzymes XbaI and SacI respectively. The *NTF* fragment was ligated into the pMDC163 Gateway destination vector (Curtis and Grossniklaus, 2003), replacing the original GUS gene, and combined with *p*HKT1 in an LR reaction. The resulting *pHKT1::NTF* construct was sequenced and inserted by heat shock into *Agrobacterium tumefaciens* strain GV3101 pMP90. *A. thaliana* Col-0 *pACT2::BirA* plants constitutively expressing biotin ligase (Deal and Henikoff, 2010) were transformed by *Agrobacterium*-mediated floral dip with the *pHKT1::NTF* construct and transgenic plants were selected for hygromycin resistance. NTF biotinylation by BirA ligase activity was confirmed by Western blotting using streptavidin-horseradish peroxidase conjugate (Thermo Scientific). The location of NTF in the nuclei of root cells was assessed through the visualisation of GFP by confocal microscopy. A homozygous *pHKT1::NTF pACT2::BirA* line identified from T4 seeds was used for the experiments (‘HKT1-INTACT line’).

### Confocal microscopy for GFP localization

A LSM 510 confocal microscope (Carl Zeiss, Jena, Germany) was used to visualise GFP fluorescence in roots after staining of cell walls with propidium iodide (10 μg/mL propidium iodide for 10 min). The excitation wavelength for both GFP and propidium iodide was 488 nm. Fluorescent signals were collected through 505-530 nm and 560-615 nm filters, respectively. Z-stacks were taken at 1 μm intervals and used to re-construct transverse cross sections with Image J.

### Plant materials

Cell-type analysis was carried out with the HKT1-INTACT line generated as part of this project (see above). Mutant analysis used *A. thaliana* knockout lines for ZAT6 and REF6, namely *ref6-3* (SAIL 747 A07), *ref6-5* (SAIL 428 A01), *zat6-1* (SALK 061991C) and *zat6-2* (SALK 050196). Seeds were germinated on vertical agar plates (10 seeds per plate) containing a minimal nutrient solution supplemented with 3% sucrose (Sani et al., 2013). Seedlings were grown for two weeks at 22°C in a 10 h light (120 μmol)/14 h dark photoperiod. At six-leaf stage, root tissue was harvested and flash-frozen in liquid nitrogen and stored at -80 ºC. Experiments were carried out with roots from five independently grown batches, each comprising approximately 200 plants (for FACS) or 1600 plants (for INTACT ChIP). For RT-qPCR three independent plant batches comprising 15 plants each were used as replicates.

### Fluorescence activated cell sorting

Protoplast generation and FACS was carried out as previously reported (Walker et al., 2017). Roots were cut into 2-3mm sections, added to 5 mL filter-sterilised solution A “plus” (0.5 M mannitol, 2 mM MgCl2, 2 mM CaCl2, 10 mM KCl, 2 mM MES hydrate, 0.1% BSA, 1.2% cellulase RS (Melford), 1.5% cellulase R10 (Duchefa), 0.2% macerozyme R10 (Duchefa) and 0.12% pectinase (Sigma), pH 5.7) and agitated at room temperature for 1 h. The sample was passed through a 70 μm filter before centrifugation at 260 rcf for 10 minutes at 4 ºC. Supernatant was removed, protoplasts were washed with 1 mL solution A (0.5 M mannitol, 2 mM MgCl2, 2 mM CaCl2, 10 mM KCl, 2 mM MES hydrate and 0.1% BSA, pH 5.7), and centrifuged again at 260 rcf for 10 minutes at 4 ºC, then resuspended in 500 μL solution A. Protoplasts were incubated on ice before FACS. A BD Influx cell sorter fitted with a 100 μm nozzle and using FACS-Flow (BD Biosciences) as sheath fluid was set to dual-gate sort (GFP, non-GFP) using a 488 nm laser. Cells were sorted into 1.5 ml buffer RLT from RNeasy Plant Mini-Kit (Qiagen) containing 1% v/v β-mercaptoethanol. RNA was extracted using this kit following manufacturer’s instructions.

### RNA sequencing

RNA-sequencing was carried out at Glasgow Polyomics (University of Glasgow). Libraries were created using Illumina TruSeq stranded mRNA kit according to the standard protocol and subsequently sequenced on Illumina NextSeq 500 sequencer using high output flow-cells to produce single-end 75 bp long reads.

### RNA-seq data analysis

The raw fasta files were pre-processed to trim the 3’ end adapter and remove low quality 3’ bases (with a PHRED score less than 15) with *cutadapt* (version 1.5) (Martin, 2011). Reads trimmed to less than 54 bases in length were also removed. Transcript expression quantification was performed using *kallisto* software (version 0.43.0) (Bray et al., 2016) against the TAIR10 transcriptome. Read counts related TAIR10 transcripts were collected into gene specific read counts. These were processed with DESeq2 software (version 1.24.0) (Love et al., 2014) to identify differentially expressed genes. Chloroplast and mitochondrial genes, and all genes with an overall base mean <1.5, were excluded from further analysis, resulting in a total number of 25144 ‘root expressed’ nuclear genes. For identification of genes with preferential expression in the pHKT-NTF tagged (T) versus non-tagged (NT) cell types we applied a cut-off of T/NT > 4-fold and p < 10^−4^. Annotation enrichment analysis was performed using DAVID (Huang et al., 2009).

### INTACT nuclei purification

Biotin-labelled nuclei purification was based on the INTACT method (Deal and Henikoff, 2010; Deal and Henikoff, 2011; Moreno-Romero et al., 2017). In short, 1 g of frozen roots were ground in liquid nitrogen and homogenised in extraction buffer (400 mM sucrose, 10 mM Tris-HCl pH 8, 10 mM MgCl2, 0.5% (v/v) Triton X-100, 5 mM β-mercaptoethanol, 1 mM PMSF and Complete protease inhibitors (Roche)). Extracts were filtered through miracloth and a 40-μm strainer. Samples were centrifuged and the pellet was resuspended in PBSb buffer (137 mM NaCl, 2.7 mM KCl, 4.3 mM Na2HPO4, 1.4 mM KH2PO4, 5 mM β -mercaptoethanol, 1 mM PMSF and Complete protease inhibitors, supplemented with 1% (w/v) biotin-free BSA (Sigma)). A fraction was kept aside on ice until further processing (‘Whole Root’ sample). The remaining fraction (‘HKT1’ sample) was incubated with PBSb-equilibrated M-280 streptavidin-coated Dynabeads (Invitrogen), washed in PBSbt buffer (PBSb with 0.1% Triton X-100) and run twice through a column-magnet system to capture the bead-bound nuclei which were re-suspended in PBS buffer, collected in a magnetic rack and re-suspended in PBS buffer. Enrichment of HKT1 nuclei was assessed by the ‘spike-in’ method (Moreno-Romero et al., 2017) and by confocal microscopy.

### Chromatin immunoprecipitation

Immediately after nuclei purification, formaldehyde was added to both ‘Whole Root’ and ‘HKT1’ samples to a final concentration of 1% (v/v) for cross-linking on ice. After quenching nuclei from the ‘Whole Root’ sample were pelleted, while bead-bound nuclei from the ‘HKT1’ sample were captured with a magnetic rack. Chromatin was sonicated to a size of approx. 400 bp and fraction kept as ‘Input’. Antibodies against H3 (Abcam, #ab1791), H3K4me3 (Diagenode, #C15410003) or H3K27me3 (Diagenode, #C15410069) were added for overnight immunoprecipitation. The chromatin-antibody complex was recovered with protein A Dynabeads (Invitrogen) and washed. DNA was eluted, reverse cross-linked and treated with proteinase K. DNA was purified using MinElute (Qiagen). Successful ChIP was confirmed through a quality control as reported before (Sani et al., 2013). Enrichment of sequences known to be associated with H3K27me3 (positive control, At5g56920) or not (negative control, At5g56900) was determined by qPCR using the Input sample for normalization. Primers used are listed in Supplemental Table 1.

### ChIP-sequencing

Sequencing of the ChIP DNA was carried out in the Glasgow Polyomics facility (University of Glasgow). DNA libraries were prepared using the the NEBNext® Ultra™ DNA Prep Kit (New England BioLabs® Inc.) according to the manufacturer’s protocols, size selected with SPRIselect Beads and amplified by PCR. The libraries were then sequenced with Illumina NextSeq® 500 system using high output flow-cells to produce single-end 75 bp long reads.

### ChIP-seq data analysis

The raw fasta files were preprocessed using *cutadapt* (version 1.5) (Martin, 2011) and *sickle* (version 0.940; flags –q 10, -l 54)(Joshi and Fass, 2011) software. Reads were aligned to the *A*.*thaliana* Col-0 genome (TAIR10) using *bowtie* (version 0.12.7) (Langmead et al., 2009). The alignment files in SAM/BAM format were sorted and duplicated reads of the same orientation removed using *samtools* (version 0.1.19) (Li et al., 2009). The resulting alignment positions were stored in BED files. For each sample aligned reads were counted in shifting 200-bp windows using *sicer* (version 1.03) (Zang et al., 2009). The profiles were stored in WIG files. For quantitative comparison of H3K27me3 we calculated for each gene the sum of read counts over seven consecutive 200-bp windows, including the TSS-containing window, one upstream and five downstream windows. Pairwise ratios of these ‘cumulative read counts’ for all replicates were analysed with the Rank Product (RP) method (Breitling et al., 2004) implemented in RankProd R (Hong et al., 2006) after applying regularized log transformation (*rlog*) within *DESeq2* R module (Love et al., 2014). Comparisons (HKT1/Whole Root) were performed twice, including or excluding two samples with low total read number. In both cases the same genes were found to show significant differences between the cell types. The p-values provided are based on all samples (6 replicates for HKT1 and 5 replicates for Whole Root) applying a cut-off at FDR < 0.05 and a rlog-based fold-change of ≥ 1.2. Annotation enrichment analysis was performed using DAVID (Huang et al., 2009).

### Quantitative PCR

qPCR was carried out on StepOne Plus (Applied Biosystems), using Brilliant III UltraFast SYBR QPCR Master Mix (Agilent). Primers are listed in Supplemental Table 2. For RT-qPCR, YSL8 (At1g48370) was used as reference. Primers for ChIP qPCR were designed to amplify two regions in ZAT6 (At5g04340) and a reference region (At5g04410). Values were normalised to Input.

### Measurement of shoot ion content

For analysis of shoot ion contents, plants were transferred from plates to hydroponics at 20 days after germination and grown on control media for another week. 100 mM NaCl was supplied (or not, control) in independent treatments of each plant. Whole rosettes were harvested 120 hours after treatment. Tissues were dried at 60 °C and dry weights determined before incubation in 5 ml 1 M HCl for 48 hours. Supernatants of extract were diluted (20-100x), and Na and K concentrations determined in a flame photometer (410, Sherwood-Scientific Ltd, UK) based on NaCl or KCl standard curves. Concentrations were calculated back to undiluted extract and normalized to shoot dry weight of each plant.

## Data availability

All data from ChIP-seq and RNA-seq are available from the European Nucleotide Archive under accession ID PRJEB39290.

## AUTHOR CONTRIBUTIONS

MAA generated the stable homozygous HKT1-INTACT plants and carried out the INTACT ChIP and ChIP-seq experiments. EMA carried out the FACS experiments together with LW, as well as ChIP-qPCR, RT-qPCR on ZAT6 and ion measurement. GP helped with the generation of the Gateway constructs and mutant lines. GH analysed the RNA-Seq data. PH supervised the sequencing and analysed the ChIP-Seq data. MLG supervised the FACS experiments at the University of Warwick. AA conceived the project, supervised all experimental work at Glasgow, and wrote the paper with assistance from all co-authors. All authors have read and approved the final manuscript version.

## Acknowledgments

We are grateful to Jordi Moreno-Romero (Uppsala University, CRAG Barcelona) for advice on the INTACT experiments, to Amparo Ruiz-Prado (Glasgow University) for horticulture assistance, to Julie Galbraith (Glasgow Polyomics) for preparing libraries and carrying out sequencing, and to Mike Blatt (Glasgow University) for help with confocal microscopy. We thank Steve Henikoff (Fred Hutch Seattle) and François Roudier (ENS, Paris, Lyon) for supplying the original INTACT constructs and lines, and Steven Jacobsen (UCLA) and José Gutiérrez-Marcos (Warwick University) for providing *ref6* seeds.

---

**Supplemental Table 1.**
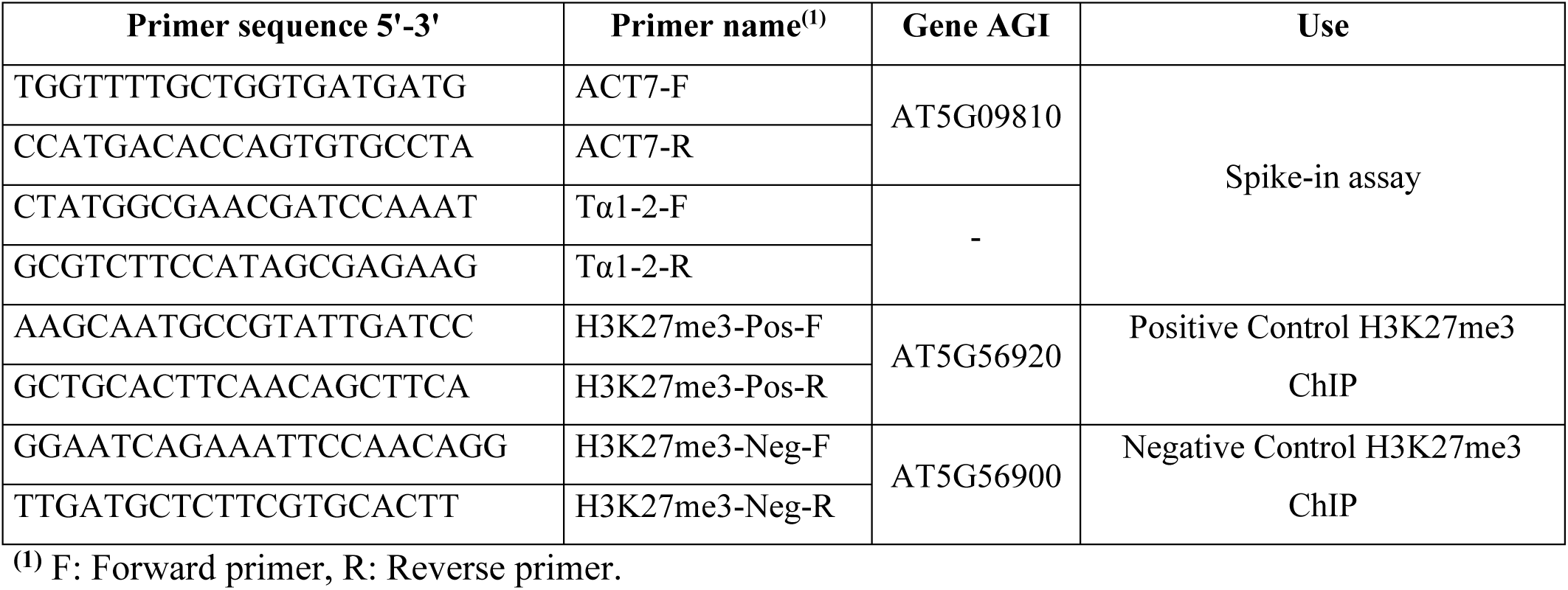
Sequences of primers used for INTACT and ChIP quality controls.

**Supplemental Table 2.**
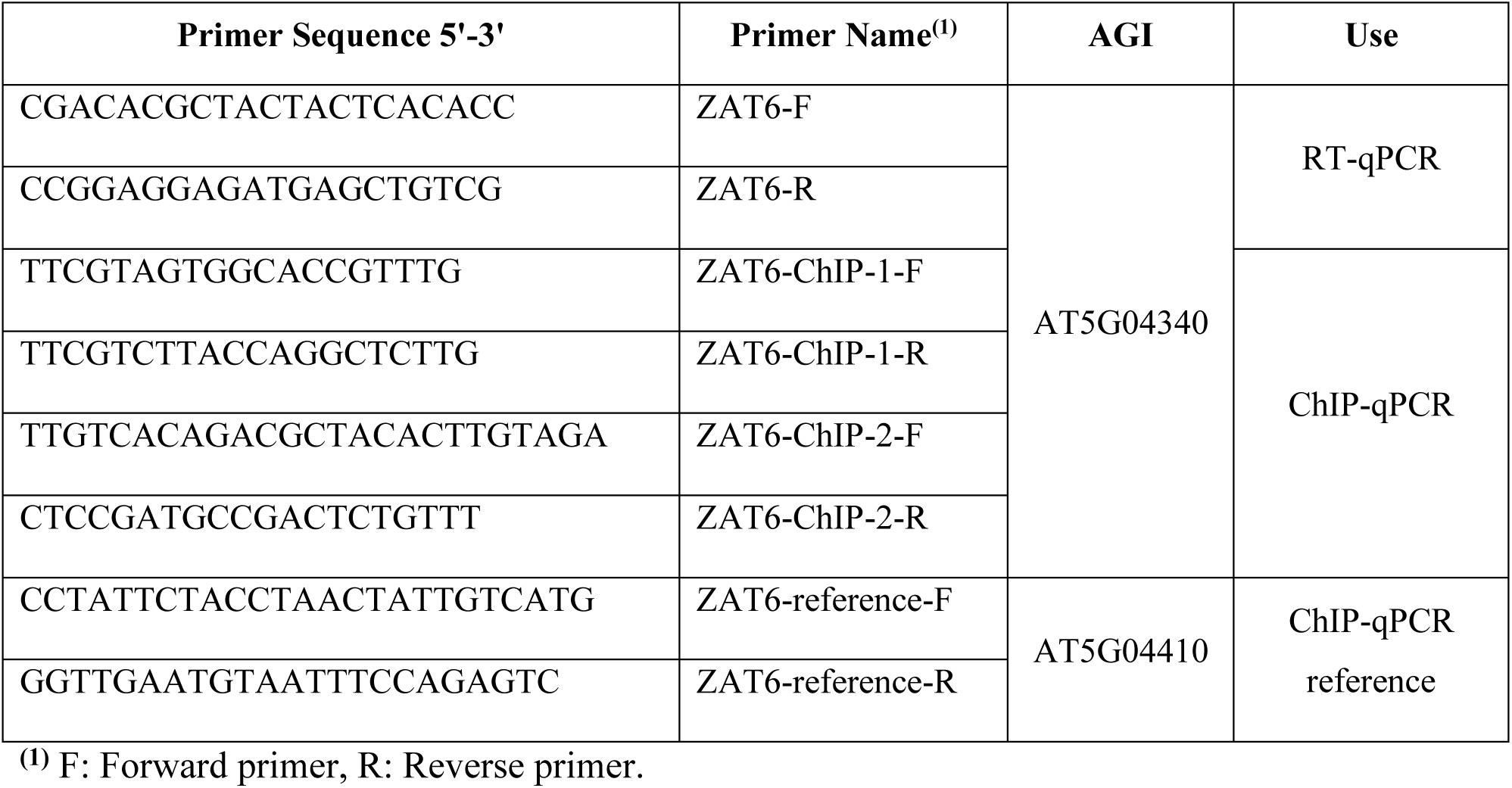
Primers used for ZAT6 ChIP-qPCR and RT-qPCR.

**Supplemental Figure 1.**
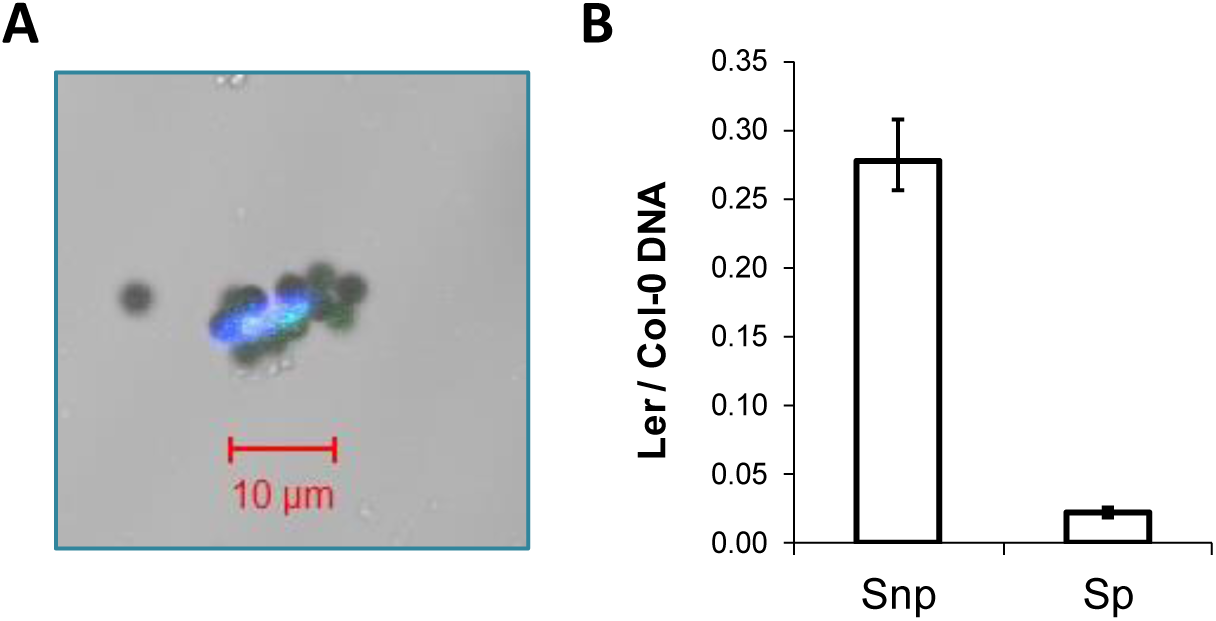
INTACT nuclei purification and determination of sample purity by the ‘spike-in’ method. **A:** Visualisation of a pHKT1-NTF tagged nucleus captured with streptavidin-coated magnetic beads under the confocal microscope. Merged image of bright field and fluorescent signals for DAPI (blue staining of nuclei) and GFP (green signal of nuclear envelope). Scale bar is 10 μm. **B:** Relative ratio of Ler to Col-0 DNA as determined by qPCR using the ‘spike-in’ method (primers listed in Supplemental Table 1). Snp: spike-in non-purified sample; Sp: spike-in purified sample. Bars represent means and standard errors of all the purifications carried out for this study (n=8).

**Supplemental Figure 2.**
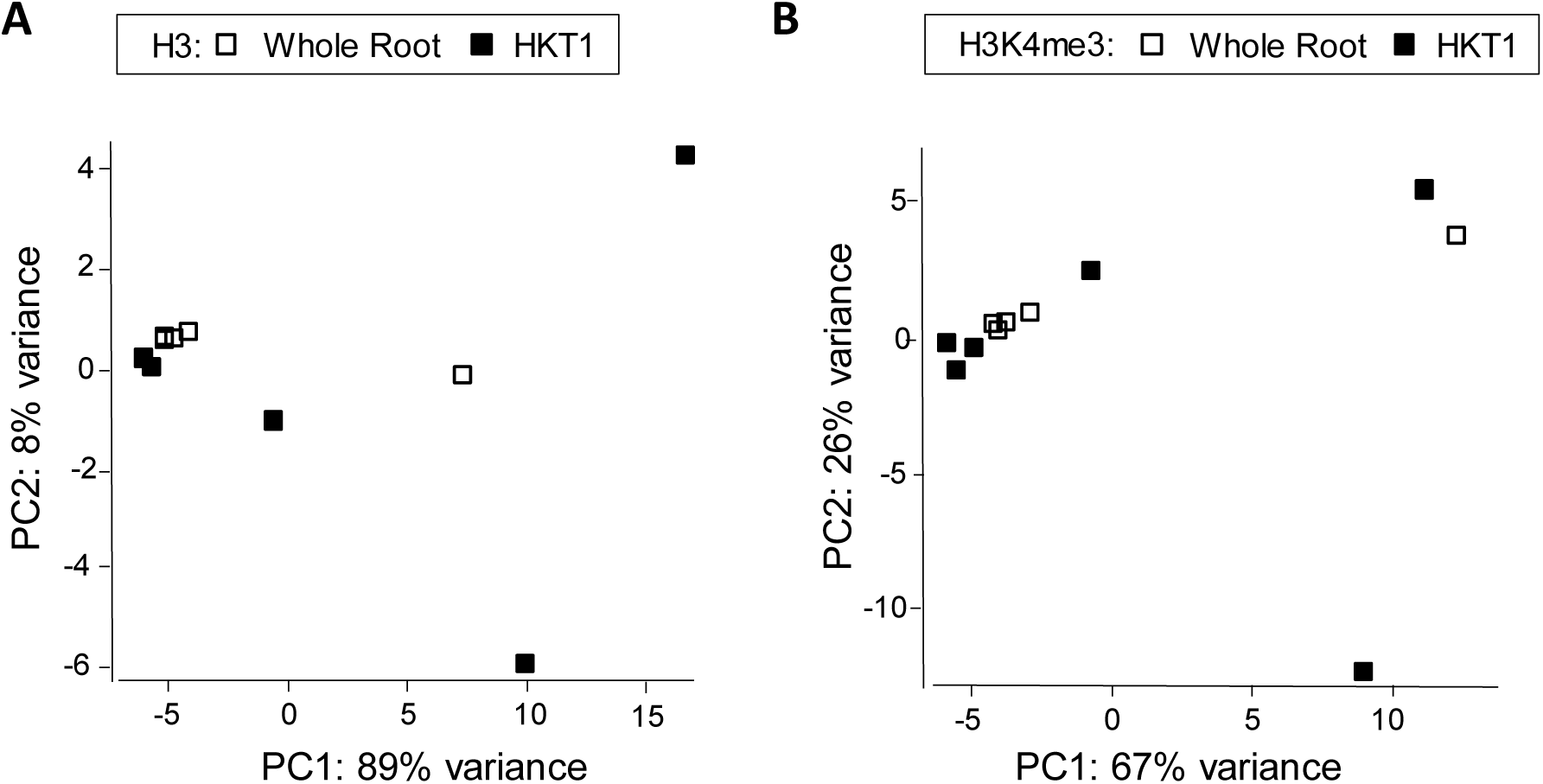
Principal component analysis based on genome wide H3 (A) and H3K4me3 (B) levels. Biotin-labelled nuclei (HKT1, black symbols) were isolated from roots of *pHKT1::NTF pACT2::BirA* lines using pulldown with streptavidin (INTACT). Total nuclear preparations (not subjected to INTACT) from the same root material represent all cell types (Whole Root, open symbols). The nuclear isolates were subjected to ChIP with antibodies against H3K27me3. Replicate samples were obtained from independently grown plant batches.

**Supplemental Figure 3.**
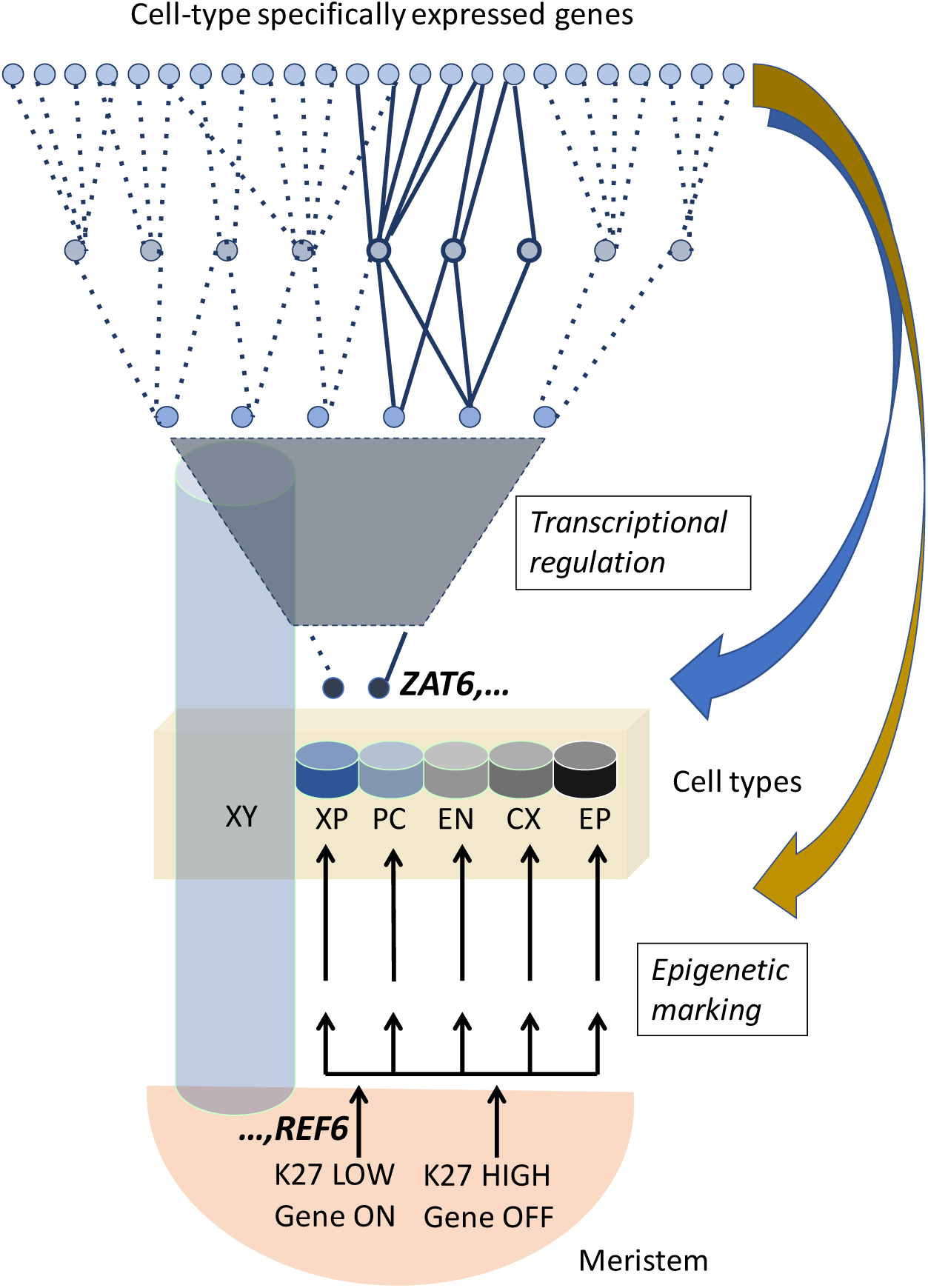
Working model for the establishment of cell-type specific transcriptional networks. In this model a relatively small number of core regulators such as *ZAT6* are cell-type specifically de-repressed during differentiation through the activity of H3K27me3 demethylases such as REF6 (K27 indicates H3K27me3). Regulation of multiple genes by each core regulators (e.g. solid lines for ZAT6-dependent genes) results in downstream proliferation of the expression pattern and establishes cell-type specific regulatory networks. The model proposes a primary mechanism for cell-type specific de-repression, which could be further enhanced through feedback regulation at the epigenetic level (brown arrow) and the transcriptional level (blue arrow), involving for example cell-type specific expression of chromatin modifiers, generation of cell-type specific signals and positional effects. Root tissues (meristem and mature root) are shown in shades of brown. Different cell types are shown in shades of blue and grey (XY: xylem, XP: xylem parenchyma, PC: pericycle, EN: endodermis, CX: cortex. EP: epidermis). For simplicity, some cell types (e.g. phloem and companion cells) are not shown here.

